# A resource for development and comparison of multi-modal brain 3T MRI harmonisation approaches

**DOI:** 10.1101/2023.06.16.545260

**Authors:** S. Warrington, A. Ntata, O. Mougin, J. Campbell, A. Torchi, M. Craig, F. Alfaro-Almagro, K. L. Miller, P. S. Morgan, M. Jenkinson, S. N. Sotiropoulos

**Affiliations:** Sir Peter Mansfield Imaging Centre, School of Medicine, University of Nottingham, UK; Sir Peter Mansfield Imaging Centre, School of Physics, University of Nottingham, UK; Wellcome Centre for Integrative Neuroimaging, FMRIB Centre, University of Oxford, UK; Australian Institute for Machine Learning, University of Adelaide, Australia; South Australian Health and Medical Research Institute (SAHMRI), Adelaide, Australia; National Institute for Health Research (NIHR) Nottingham Biomedical Research Centre, Nottingham, UK

**Keywords:** travelling heads, between-scanner, open data, COMBAT

## Abstract

Despite the huge potential of magnetic resonance imaging (MRI) in mapping and exploring the brain, MRI measures can often be limited in their consistency, reproducibility and accuracy which subsequently restricts their quantifiability. Nuisance nonbiological factors, such as hardware, software, calibration differences between scanners, and post-processing options can contribute to, or drive trends in, neuroimaging features to an extent that interferes with biological variability. Such lack of consistency, known as lack of harmonisation, across neuroimaging datasets poses a great challenge for our capabilities in quantitative MRI. Here, we build a new resource for comprehensively mapping the extent of the problem and objectively evaluating neuroimaging harmonisation approaches. We use a travelling-heads paradigm consisting of multimodal MRI data of 10 travelling subjects, each scanned at 5 different sites on 6 different 3T scanners from all the 3 major vendors and using 5 neuroimaging modalities, providing more comprehensive coverage than before. We also acquire multiple within-scanner repeats for a subset of subjects, setting baselines for multi-modal scan-rescan variability. Having extracted hundreds of image-derived features, we compare three forms of variability: (i) between-scanner, (ii) within-scanner (within-subject), and (iii) biological (between-subject). We characterise the reliability of features across scanners and use our resource as a testbed to enable new investigations that until now have been relatively unexplored. Specifically, we identify optimal pipeline processing steps that minimise between-scanner variability in extracted features (implicit harmonisation). We also test the performance of post-processing harmonisation tools (explicit harmonisation) and specifically check their efficiency in reducing between-scanner variability against baseline standards provided by our data. Our explorations allow us to come up with good practice suggestions on processing steps and sets of features where results are more consistent, while our publicly-released datasets establish references for future studies in this field.

## 1. Introduction

A key challenge in extracting robust quantitative information from magnetic resonance imaging (MRI) data of the brain is the dependence of imaging-derived phenotypes (IDPs) on nuisance non-biological factors. These factors range from hardware and software differences, and scanning protocol parameters and implementation, which are different between vendors and can vary with site (Han et al., 2006; Zhu et al., 2011) and following scanner upgrades (Jovicich et al., 2009; Potvin et al., 2019). Additionally, image processing options (for IDP extraction for example) vary across research groups, thus introducing additional non-biological sources of variability. Such factors can affect IDPs in non-trivial ways (Takao et al., 2011; Zhu et al., 2011), leading to biases and increased variability in measurements obtained from different settings (Chen et al., 2014; Jovicich et al., 2006; Vollmar et al., 2010). This is true, even in cases where scans have been acquired with a rigid acquisition protocol or calibrated with phantoms; quantitative measurements can still show variance reflecting non-biological causes (Cheng and Halchenko, 2020; Lee et al., 2021).

This lack of consistency or “*harmonisation”* across sites and scanners impedes and reduces the potential for quantitative applications of MRI. At the extreme, variability of measures obtained from the same subject but on different scanners can be as large as biological between-subject variability (Mirzaalian et al., 2016), creating obvious interpretation issues and questions on usefulness of some of these metrics in real-world scenarios (Rao et al., 2017). Reduced quantifiability can have downstream effects on the reproducibility and generalisability of findings and direct consequence in two key scenarios: (i) the pooling of multi-site neuroimaging datasets (Zhu et al., 2011), potentially acquired at also different times, and (ii) relating new IDPs acquired under different scanning conditions to an existing set of normative data (Bayer et al., 2022). The pooling of multi-site neuroimaging datasets is arguably the most sustainable way for having studies of larger scale and for increasing the diversity of cohort demographics, a key factor in ensuring robust and generalisable science (Oh et al., 2015). Non-biological sources of variation have a direct negative effect on this pooling, and thus on reproducibility and representation of diverse populations. The construction of normative models, i.e. models that capture healthy biological variation of a phenotype (Marquand et al., 2016), is vital in the uptake of quantitative MRI in the clinical setting. Non-biological sources of variation hinder comparisons of newly acquired data with normative models, reducing the confidence in whether deviations are due to biological or non-biological effects. Hence, visual (and therefore subjective) inspection by radiologists is still the preferred way forward in clinical settings. Finally, non-biological variability may mask true effects, or lead to the false conclusion of group differences, thereby not only affecting reproducibility, but also the sensitivity and specificity of a study. Such challenges also underlie the relatively limited, albeit growing, uptake/success of modern MRI technologies in clinical trials (Ellingson et al., 2015; Sadraee et al., 2021; Nigri et al., 2022).

A good preliminary step for minimising these issues is to ensure the standardisation of scanning protocols across scanners/sites (Chalavi et al., 2012). However, this is a non-trivial task that is not always scalable or practical and does not resolve the problem fully. Firstly, vendor-specific proprietary implementations can often lead to only nominal matching of parameter acquisitions rather than true matching, causing signal/contrast/distortion differences. Secondly, expert knowledge of these implementation differences by local physicists are needed, which is not always available. Thirdly, even using the same raw datasets acquired using the same protocols, variability in processing and filtering options can lead to significantly different IDPs and results (Griffanti et al., 2016; Botvinik-Nezer et al., 2020; Schilling et al., 2021). Harmonisation therefore needs to be considered at all points of a study, from design and data acquisition, to data processing and IDP extraction. Attempts to standardise acquisition alone will most likely lead to *aligned* protocols, but with inevitable differences across platforms.

For that reason, post-acquisition harmonisation approaches of neuroimaging data have been developed (Fortin et al., 2018; Cetin Karayumak et al., 2019) that aim to remove non-biological variability while still preserving variance in IDPs associated with biological factors. Such approaches are likely to have higher success rates when some effort is first made to align acquisition protocols. In general, harmonisation methods fall into two main categories, depending on whether they harmonise IDPs directly (Fortin et al., 2018; Yamashita et al., 2019; Garcia-Dias et al., 2020) or indirectly, by standardising the raw scans (Cetin Karayumak et al., 2019; Mirzaalian et al., 2016; Tax et al., 2019). Nevertheless, what is generally missing are objective ways and datasets to evaluate and compare such approaches. Different studies have relied so far on a range of indirect metrics, from using population distributions as a reference (Garcia-Dias et al., 2020), to subject group matching by attributes such as age, sex, gender, race and handedness (Fortin et al., 2017). An alternative and more direct approach for assessing the quality of harmonisation is to use within-scanner repeats. For example, in (Vollmar et al., 2010), two within-scanner repeats were used as a baseline within-subject variability reference towards which harmonisation success was assessed. In (Kurokawa et al., 2021), two within-scanner repeat scans from four subjects, and scan-rescan Human Connectome Project (HCP) (Van Essen et al., 2013), data were used as a baseline. Despite these previous efforts, there is still limited understanding of which brain MRI modalities and which IDPs within each modality are less sensitive to between-scanner effects and hence will benefit less/more from harmonisation methods.

In this study, we provide a resource aimed at better understanding the nature of the challenge and for setting the foundations to address it. Firstly, we present a unique comprehensive dataset for multi-modal brain MRI harmonisation acquired using a travelling-heads paradigm; ten healthy individuals scanned multiple times across multiple sites and scanners using T1-weighted (T1w), T2-weighted (T2w), susceptibility-weighted (SWI), diffusion MRI (dMRI) and resting-state functional MRI (rfMRI) sequences. We extend previous similar approaches (Pohl et al., 2016; Tax et al., 2019; Yamashita et al., 2019; Tong et al., 2020; Maikusa et al., 2021; Kurokawa et al., 2021; Tanaka et al., 2021; Duff et al., 2022; Tian et al., 2022) in a number of ways: (i) by considering scanners from all three major vendors, (ii) by considering multiple generations of scanners within each vendor, (iii) by having multiple within-scanner repeats for the same subjects, (iv) by acquiring multiple neuroimaging modalities and (v) by collecting data at five imaging sites in total. We use the UK Biobank imaging protocol (Miller et al., 2016) as a rough guide to align protocols, but within that scope we intentionally avoid nominal matching of acquisition parameters and allow for reasonable variation. This approach enables us to reflect more realistic scenarios and leverage the strengths of each considered system by preserving best practices at each imaging site.

Subsequently, we use this data resource to map the extent of the problem in hundreds of IDPs. For each of these IDPs, we compare between-scanner variability against within-scanner variability, as well as biological variability, and also explore the consistency of cross-subject ranking across scanners. We further demonstrate how we can evaluate existing harmonisation approaches, such as ComBat and CovBat (Fortin et al., 2017, 2018; Chen et al., 2022) (*explicit harmonisation*), as well as comparing the robustness and precision of image processing pipeline alternatives in extracting specific IDPs when handling data from multiple scanners (*implicit harmonisation*). We find that implicit harmonisation can offer complementary benefits to explicit harmonisation in the explored examples; and that between-scanner reliability of very commonly-used IDPs, such as cortical or subcortical volumes, can be significantly affected by how data is handled and processed. We also showcase how the acquired within-subject, within-scanner repeats can highlight challenges for existing filtering algorithms (such as diffusion MRI denoising (Veraart et al., 2016)), stemming from non-linear effects that appear to be common across different scanners, contrary to expectation. The data is publicly released in BIDS format via OpenNeuro and will be further augmented with more scanners and subjects in the near future. In addition, we make the processing pipeline and resultant IDPs available.

## 2. Methods

### 2.1 Data Acquisition

We used a travelling-heads paradigm to acquire multimodal brain MRI data of 10 healthy travelling subjects (two females, eight males, age range: 24-48), each scanned on six different 3T scanners covering all three major vendors (Siemens/Philips/GE), from five different sites, and covering a range of hardware features (for instance bore size, gradient strength, number of head coil channels, acceleration capabilities). For a subset of four subjects we acquired five additional within-scanner repeats using a different scanner for each subject (i.e. for each subject we had six within-scanner sessions for one scanner and one session on the remaining five scanners), resulting in 80 sessions in total. In each session, five imaging modalities were acquired: T1w, T2w, SWI, dMRI, and rfMRI. Scanner details are summarised in Figure 1 and subject demographics are summarised in Supplementary Table 1. The within-scanner repeats were acquired using the Philips Achieva, Siemens Prisma (32ch), Siemens Trio and Siemens Prisma (64ch) systems.

**Figure 1.**
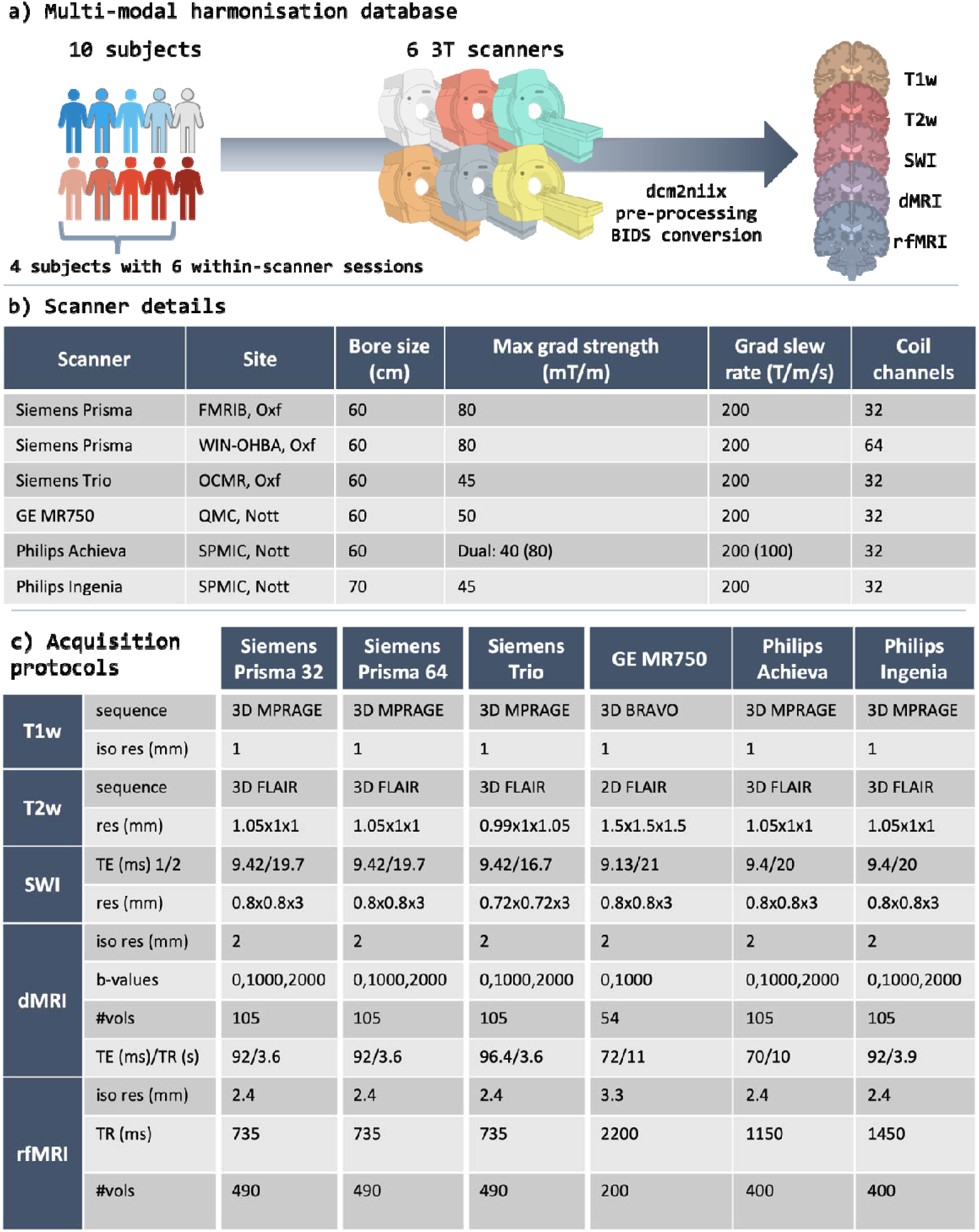
The multi-modal harmonisation database. a) Ten subjects were scanned on six 3T scanners covering the three main vendors (GE, Siemens, Philips) at five different sites. On four scanners, one of the subjects was chosen to complete five within-scanner repeats. In each session five modalities are acquired: T1-weighted, T2-weighted, susceptibility-weighted imaging, diffusion MRI, and resting-state rfMRI. Data were pre-processed and converted to BIDS format, which are publicly available. b) A summary of key scanner details and specifications. Oxf = University of Oxford; Nott = University of Nottingham. c) A summary of key acquisition parameters for the five modalities, for all six scanners, highlighting parameters that vary across scanners. A blip-reversed spin-echo fieldmap was also acquired for correcting susceptibility-induced distortions with the phase encoding direction switching along the anterior-posterior orientation.

Data acquisition was performed under two ethics protocols for healthy volunteers at Nottingham (PI: Sotiropoulos, Ethics: FMHS-36-1220-03, H14082014/47). The Oxford data acquisition was performed under an agreed technical development protocol approved by the Oxford University Clinical Trials and Research Governance office, in accordance with International Electrotechnical Commission and United Kingdom Health Protection Agency guidelines. Informed consent was obtained from all participants.

Scanning protocols were *guided* by the UK Biobank (UKBB) neuroimaging study (Miller et al., 2016), which is a relatively short multi-modal protocol (about 35 minutes in total), that does not rely heavily to specialised hardware/software and hence it is anticipated to be relatively generalisable across scanners. We did not aim to perfectly match every single parameter in this protocol, but instead respected best practice for each scanner/site and remained within the limitations of scanner hardware/software. Perfectly matching protocols is not always possible, nor realistic; and it can lead to nominal-only matching of acquisition parameters, rather than matching of image quality and features across scanners. We show in the Supplementary Material (Supplementary Figure 1, and discussion below) an example case for resting-state functional MRI. Protocol summaries are provided in Figure 1, highlighting differences between scanners. Shimming was performed at the beginning of each session and auto-reshimming was disabled. To correct for susceptibility-induced distortions for dMRI and rfMRI, we acquired a blip-reversed spin-echo fieldmap (Andersson et al., 2003) with the phase-encoding (PE) direction switching along the anterior-posterior orientation. The same PE direction was used for dMRI and rfMRI in each session.

#### T1-weighted

We used T1w gradient echo (3D MPRAGE (Mugler III and Brookeman, 1990) for Siemens and Philips scanners, 3D BRAVO for the GE MR750) scans with an isotropic spatial resolution of (1mm)^3^. As in the original UKBB protocols, gradient non-linearity distortion correction (GDC) was turned off for the Siemens scanners because the Siemens on-scanner corrections have been found to provide inconsistent results, particularly for 2D EPI acquisitions (scanner-corrected 3D and 2D acquisitions of the same subject cannot be successfully aligned with a rigid body transformation). Instead, these corrections were performed offline using vendor-supplied gradient non-linearity descriptor files (Alfaro-Almagro et al., 2018). For the non-Siemens scanners, GDC correction was performed on the scanner. This applies to all other modalities we acquired. Vendor-provided pre-scan normalise was used for all scanners. Scan time was on the order of five minutes.

#### T2-weighted FLAIR

With the exception of the GE MR750, all the T2w scans were performed using a 3D T2w FLAIR sequence that allowed high-resolution data (almost (1mm)^3^ isotropic) in four minutes. The software version on the MR750 did not have 3D T2w FLAIR functionality (i.e., it could either provide a 3D FLAIR with no T2-weighting or a 2D T2w FLAIR). Therefore, we obtained a 3D FLAIR without T2w and also acquired a 2D T2w FLAIR, which is inherently slower than 3D and compromised with spatial resolution. We acquired three versions: (i) 1mm isotropic 3D FLAIR, (ii) 1.5mm isotropic 2D T2w FLAIR, and (iii) 1×1×2mm 2D T2w FLAIR. The same GDC and pre-scan normalise options were followed as before. For analysis we used the 1.5mm isotropic 2D FLAIR for the GE scans, but we provide the others as well in the public release.

#### Susceptibility-weighted imaging (SWI)

The SWIs were acquired using anisotropic, complex data for two echoes, roughly matching around TE_1_∼9s and TE_2_∼20s. For the GE scanner we used the SWAN sequence, which acquired seven echoes, and the two echoes closer to TE_1_ and TE_2_ were extracted during processing. This resulted in a higher bandwidth for the GE data (∼350 Hz/pixel for GE vs ∼140 Hz/pixel for Philips and Siemens). Accurate reconstruction of phase images required the complex sensitivity of the individual coil data as anomalous phase transitions in regions of focal dropout have been reported (Alfaro-Almagro et al., 2018; Robinson et al., 2017). For the Siemens scanners, as in the original UKBB protocol, data from individual coils were saved separately, and phase images were subsequently high-pass filtered and combined during post processing. For the non-Siemens scanners, such anomalous phase transitions are less common and hence individual coil data were combined on the scanner. Magnitude and phase images were saved for all the scanners. Scan times were on the order of 2.5 minutes for all scanners.

#### Diffusion MRI (dMRI)

The diffusion images were acquired with a monopolar pulsed gradient spin echo (PGSE) EPI sequence at (2mm)^3^ isotropic spatial resolution. We used an anterior-posterior phase encoding direction and acquired reversed spin-echo EPI b=0 s/mm^2^ scans on all scanners. Differences in gradient strength and simultaneous-multi-slice (multiband) acceleration capabilities affected the achievable minimum TE and TR across scanners. Both the Philips Achieva and GE MR750 did not have multiband capabilities, therefore the resulting TR was above 10 seconds. For the MR750, we opted for only relatively low b-value data (up to b=1000 s/mm^2^), because of the low gradient strength and excessively long TR. TR was also long for the Philips Achieva, but the much stronger gradients allowed usable data in a reasonable scan time. In the absence of multi-slice acceleration for the Achieva and MR750, in-plane parallel imaging with an acceleration of two was used to minimise TE. We were able to approximately match angular resolution across b-shells for all scanners. In summary, total scan times were on the order of 6.5 minutes for the Siemens scanners, 7.5 minutes for the Philips Ingenia, 18 minutes for the Philips Achieva and 12 minutes for the GE MR750.

#### Resting-state functional MRI (rfMRI)

The rfMRI images were acquired with 2D gradient echo planar imaging (GE EPI). All subjects were asked to keep their eyes open during scanning. As in dMRI, deviations from the UK Biobank protocols were required due to the differences in the acceleration capabilities of each scanner. We acquired two sets of rfMRI data for the GE MR750 and Philips Ingenia using a) protocols that were as nominally-matched as possible and b) protocols that were more in-line with scanner-specific best practices. We compared image quality across scanners in each case. For the Philips Ingenia scanner, pushing the multiband acceleration factor beyond four caused excessive artefacts and data had reduced temporal signal to noise ratio (tSNR). In comparison, we were able to achieve a multiband acceleration factor of eight on Siemens scanners without problematic artefacts. We therefore opted for acquisitions that had the same spatial resolution as the Siemens scanners and roughly the same number of timepoints (400 in Philips vs 490 in Siemens) but differed in the temporal resolution. For GE (no multiband available), we accepted a reduced spatial resolution (3.3mm isotropic compared to 2.4mm isotropic with Siemens) in order to keep tSNR more consistent with Siemens’ data. In total the scan times were 6 minutes for Siemens scanners, 7.5 minutes for the Philips Achieva, 9.5 minutes for the Philips Ingenia and 7.5 minutes for the GE MR750. In each case, the flip angle was set to the Ernst angle for the corresponding TR, assuming T_1_=1.5 s for grey matter at 3T. A summary of fMRI data image quality metrics is provided in Supplementary Figure 1, comparing all the alternatives.

### 2.2 Data Processing

#### Imaging-Derived Phenotype Extraction

Hundreds of multi-modal IDPs were extracted from each session. First, raw data were converted to NIFTI format using dcm2niix (v1.0.20211006) (Li et al., 2016) and subsequently converted to the BIDS data structure (Gorgolewski et al., 2016), before applying an adapted version of the UKBB pipeline (Alfaro-Almagro et al., 2018). All data have been anonymised, while the high-resolution anatomical images have been “defaced”. Anonymised and defaced BIDS format data are publicly available via OpenNeuro (made available on publication). This database will be further augmented in the coming years with more subjects and scanners (in particular two GE Premier Signa wide-bore 3T scanners at two different sites).

For dMRI and rfMRI data, we obtained the effective echo spacing and total readout time required for susceptibility-induced distortion correction using spin-echo fieldmaps (Andersson et al., 2003). These were extracted from dcm2niix, which takes into account nominal echo-spacing, in-plane acceleration, as well as bandwidth and matrix dimensions. The Supplementary Information summarises the equations used by dcm2niix to calculate the total readout times and Supplementary Tables 2-3 provide a summary of acceleration factors and the associated effective echo spacings and total readout times across the scanners.

We obtained image quality metrics (IQMs) in order to characterise each of the scanning sessions. We used MRIQC (v22.0.6) (Esteban et al., 2017) for T1w (e.g. smoothing extent, SNR, tissue-specific SNR and regional CNR) and rfMRI (e.g. smoothing, tSNR, motion artefact measures) data, while for dMRI we used eddyQC (Bastiani et al., 2019) to quantify SNR, angular CNR, motion and outliers. A summary of IQMs is provided in Supplementary Table 4.

A modified version of the UKBB pipeline (Alfaro-Almagro et al., 2018) was applied to extract IDPs, providing a full processing stream for all acquired modalities, from allowing data in different formats from different vendors, distortion correction and template alignment, to generating a set of IDPs for each session and subject. The pipeline was originally designed for Siemens-acquired UKBB data. We adjusted the pipeline in various ways to allow the processing of data obtained from other vendors and modified acquisition protocols. We also augmented the pipeline to allow additional processing steps/tools. For instance, we replaced the original tractography processing with the XTRACT toolbox (Warrington et al., 2020), we replaced the approximate NODDI-AMICO fit (Daducci et al., 2015) with a GPU-accelerated NODDI model (Zhang et al., 2012) fitting routine (Hernandez-Fernandez et al., 2019), we added the option for performing dMRI denoising (Veraart et al., 2016), and added the option of gradient non-linearity distortion correction. We derived multi-modal IDPs including a range of structural, microstructural, connectional and functional IDPs, specifically: volumes of tissue types; cortical surfaces and their metrics (volumes, curvature, thickness, area); subcortical region-wise volumes; measures of white matter microstructure within various white matter tracts; iron deposition proxies in grey matter; and measures of regional functional connectivity. An overview of the IDPs extracted from each modality is shown in Supplementary Figure 2.

#### Mapping Between-Scanner Effects

The extracted IDPs and IQMs may be used to assess between-scanner effects and assess variability in data quality and IDP values across scanners. We first used the IQMs to explore the presence of any outliers either across scanners or subjects in terms of overall data quality. To do so, IQMs reflecting the image quality of the anatomical (T1w), microstructural (dMRI) and functional (rfMRI) data were (i) z-scored across scanners and averaged across subjects, providing a measure of scanner data quality relative to other scanners, and (ii) z-scored across subjects and averaged across scanners, providing a measure of subject data quality relative to other subjects. In each case, to avoid bias towards any given scanner, we excluded within-scanner repeats. We also excluded IQMs describing the *b*=2,000 s/mm^2^ dMRI data as these were not available for all scanners.

Next, we assessed the between-session IDP similarity *P*_*ij*_ to reflect how similar IDPs from sessions *i* and *j* are on average (*i, j = 1: N*_*ses*_ where *N*_*ses*_ *=* 80 in our data, spanning all subjects and scanners). IDPs were grouped into *m = 1: M*_*cat*_ categories, including subcortical volumes, brain tissue volumes, subcortical T2*, cortical parcel volumes, dMRI regional and tract-wise microstructure (FA, MD, MO, L1, L2, L3) and rfMRI functional connectivity node amplitudes and edges. For each of the *M*_*cat*_ IDP categories, the Spearman’s rank correlation was calculated between pairs of sessions *i* and *j*, giving in total *M*_*cat*_ correlation values 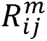, one for each IDP category. The median correlation across all IDP categories was used to reflect the between-session similarity for sessions *i* and *j*:

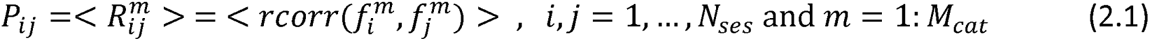

where *rcorr* is the Spearman’s rank correlation, < > is the median across *m* and 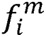 is a vector containing the IDPs for session *i* and category *m*. Note that for functional connectivity we used the IDPs extracted from a 25-dimensional group ICA with partial correlation as a connectivity measure (giving 210 edges and 21 node amplitudes). To reduce the dimensionality, we kept only the top 5% (31) strongest edges. We identified the top 5% strongest edges by calculating the mean edge weight across within-scanner repeats for each of the subjects with within-scanner repeats. The top 5% strongest edges were used throughout these analyses.

Subsequently, for each of the extracted IDPs, we calculated the coefficient of variation (CoV) across the between-scanner repeats of a subject (i.e., between-scanner, within-subject) and we compared it with two baselines: (i) the CoV of within-scanner, within-subject repeats, (ii) the CoV of within-scanner, between-subject repeats. The former provides a measure of within-scanner variability to compare against and the latter a measure of between-subject (biological) variability. We also compared IDP bias by exploring the agreement of the mean across between-scanner measurements against the mean across within-scanner measurements.

Finally, we explored how the ranking of subjects varied across scanners for each IDP *d*, i.e. quantifying the consistency 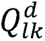 in the rank ordering of subject IDPs between scanners *l, k = 1: N*_*scan*_ (where *N*_*scan*_ *= 6* is the number of scanners and *d = 1: D* the list of all IDPs). To do so, for each IDP *d*, we calculated the Spearman’s rank *rcorr* across the ten subjects between all scanner pairs. We compared ranking consistency after grouping IDPs into sub-categories and in the case where all scanners are included (*N*_*scan*_ *= 6*) and in the case where the pool of scanners is restricted to those from a single vendor (*N*_*scan*_ *= 3* Siemens scanners). We assessed ranking consistency against an indicative “null” baseline; this was obtained by simulating random rankings, calculating the Spearman’s rank correlation, and taking the interquartile range of the distribution of correlation values.

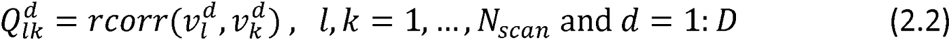

where 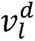 is a vector containing the IDPs for all subjects for scanner *l* and IDP *d*.

### 2.3 Evaluating Harmonisation Approaches

We utilised our data resource as a testbed for existing harmonisation approaches. Having within-scanner repeats, as well as scans of the same brain across multiple scanners, allows for multiple explicit and quantitative comparisons. As an exemplar for this study, we used the within-scanner variability as a baseline and we assessed how closely harmonisation approaches can bring between-scanner variability to this baseline for different IDPs. We also explored how stability of between-subject ranking can be affected by harmonisation approaches. We explored two groups of methods: a) implicit harmonisation: given the plethora of processing approaches for extracting the same IDPs from neuroimaging data, we evaluated how robust and consistent different approaches are in extracting the same IDPs across scanners in the same subject. We postulate that an optimal processing pipeline is as immune as possible to site/scanner effects and returns similar values for the same IDPs in the same subject scanned in various systems. We demonstrate how our database can be used for pipeline optimisation to maximise reproducibility and robustness. b) Explicit harmonisation: we used our resource to directly evaluate approaches that have been explicitly designed to remove nuisance scanner (i.e. “batch”) effects; and characterise their efficacy across different modalities and IDPs.

#### Implicit Harmonisation

First, we compared approaches for extracting subcortical volumes from anatomical images using both unimodal and multimodal subcortical segmentation. Specifically, we compared a unimodal subcortical segmentation approach (FSL’s FIRST (Patenaude et al., 2011)) to the more recently developed unimodal/multimodal FSL’s MIST (Visser et al., 2016). MIST was run in three ways: (i) using only T1w data, providing a direct comparison with FIRST, (ii) using two modalities, T1w and T2w data, and (iii) using three modalities, T1w, T2w and dMRI data. For the multimodal runs, we registered to the T2w and dMRI data to the T1w data. For the T2w registration, we used linear registration and for the dMRI data we used a boundary-based registration (Greve and Fischl, 2009). In each case MIST was trained using all sessions excluding the within-scanner repeats (60 sessions in total) and the trained model was subsequently applied to all sessions to extract subcortical segmentations. The set of subcortical structures were restricted to those available from both approaches, which includes left/right thalamus, pallidum, putamen, hippocampus, amygdala, and caudate nucleus combined with nucleus accumbens. We then compared subcortical volume variability for within- and between-scanner repeats and preservation of subject ranking across the three approaches, and against segmentations derived from FSL’s FIRST.

As a second example of pipeline optimisation, we compared approaches for deriving cortical region volumes. Specifically, we compared (i) the atlas-based approach used in the UK Biobank pipeline, where atlas-based registered ROIs are constrained by the subject-specific grey matter mask, (ii) FreeSurfer (v7.1.0) (Dale et al., 1999) and (iii) the recently developed FastSurfer (v2.0.0) (Henschel et al., 2022, 2020), a deep learning alternative to FreeSurfer. These steps provided coarse and fine resolution cortical parcellations for each subject, that were then compared.

Finally, we used a further example to demonstrate the richness of our resource in using within-scanner repeats to evaluate pre-processing steps. We assessed the effect of dMRI denoising on variability of microstructural IDPs, such as tract-wise FA and MD. As we expect thermal noise to be a large contributing factor to within-scanner, within-subject variability, we assessed whether dMRI denoising approaches reduce within-scanner variability across a range of IDPs. To do so, we denoised the raw dMRI data using MP-PCA (Veraart et al., 2016) (as implemented in MRtrix3 v3.0.2 (Tournier et al., 2019)), prior to any other processing. The denoised data were then processed using the UKBB pipeline to generate the standard dMRI IDPs. We then compared the variability of IDPs across within-scanner repeats from our pipelines run with and without denoising. In addition, we repeated the above processing but applied the denoising step after distortion corrections.

#### Explicit Harmonisation

We explored explicit harmonisation methods using our dataset. Specifically, we applied ComBat (Fortin et al., 2017) and CovBat (Chen et al., 2022) to a representative set of IDPs: atlas-based cortical grey mater volumes and subcortical volumes derived from T1w, subcortical T2* derived from SWI, and tract-wise microstructural measures (mean fractional anisotropy) derived from dMRI. We applied each harmonisation approach to the whole cohort and compared how between-scanner CoVs before and after harmonisation compares against within-scanner repeat CoVs. We also explored how harmonisation approaches affect between-scanner stability of subject ranking. For both ComBat and CovBat, subject demographics (age, sex) were used as covariates.

## 3. Results

### 3.1 A Comprehensive Multi-Modal Harmonisation Resource

In total, 80 sessions were acquired from 10 subjects (60 between-scanner and 20 within-scanner repeats). Qualitative demonstrations of the multi-modal data for a single subject across the 6 scanners are shown in Figure 2. Consistency in quality and contrast can be observed in general for all modalities/scanners, although, as expected, there are appreciable differences between scanners. Supplementary Figure 3 examples modalities where between-scanner differences are more/less appreciable. For example, dMRI-derived FA maps show greater between-scanner differences compared to within-scanner repeats. On the other hand, between-scanner variability in T1w scans are, qualitatively, comparable to within-scanner variability. These results provide an early demonstration that inter-site effects and the need for harmonisation are not equivalent across imaging modalities and IDPs.

**Figure 2.**
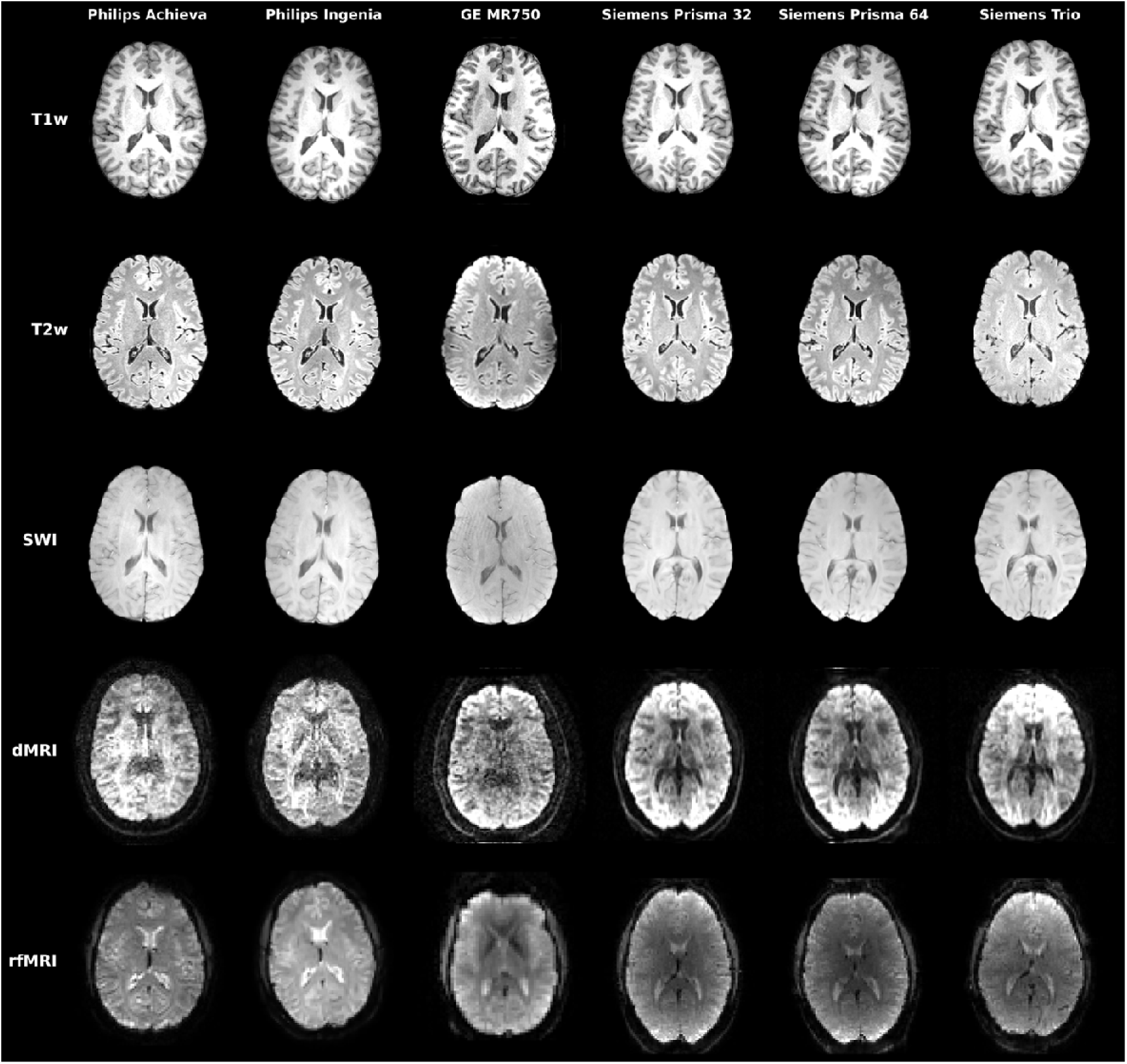
Illustration of acquired multi-modal data for a single subject across all 6 scanners and 5 imaging modalities. For dMRI, a single b=1000 s/mm^2^ is shown corresponding to the same diffusion-sensitising orientation (left-right orientation).

To perform a more quantitative comparison across scan sessions, quality control was performed, as described in Methods. The scanner/subject averaged z-scored IQMs are shown in Figure 3 for each of the considered IQMs. In the case of scanner performance (Figure 3a), since three out of six scanners were Siemens, we expect the mean IQM values to be significantly determined by the systems of this vendor. Indeed, IQMs for the Siemens scanners are closer overall to the means (i.e. z-scores closer to zero), with some modality-specific differences. Nevertheless, we observe that all metrics for all other scanners are within two standard deviations of their respective means, i.e. there are no major outliers in terms of raw image quality and/or artefacts (74% of the IQMs are within one standard deviation from their respective means). The Philips Achieva T1w and dMRI data are also closer to the mean scanner quality, while the GE rfMRI is closer to the respective rfMRI IQM mean. Similarly, at the subject-level, we find that the vast majority of IQMs (99%) are within two standard deviations from their respective means. In summary, there were no scanners/subjects in our cohort that were different enough to be considered outliers with respect to the other observations.

**Figure 3.**
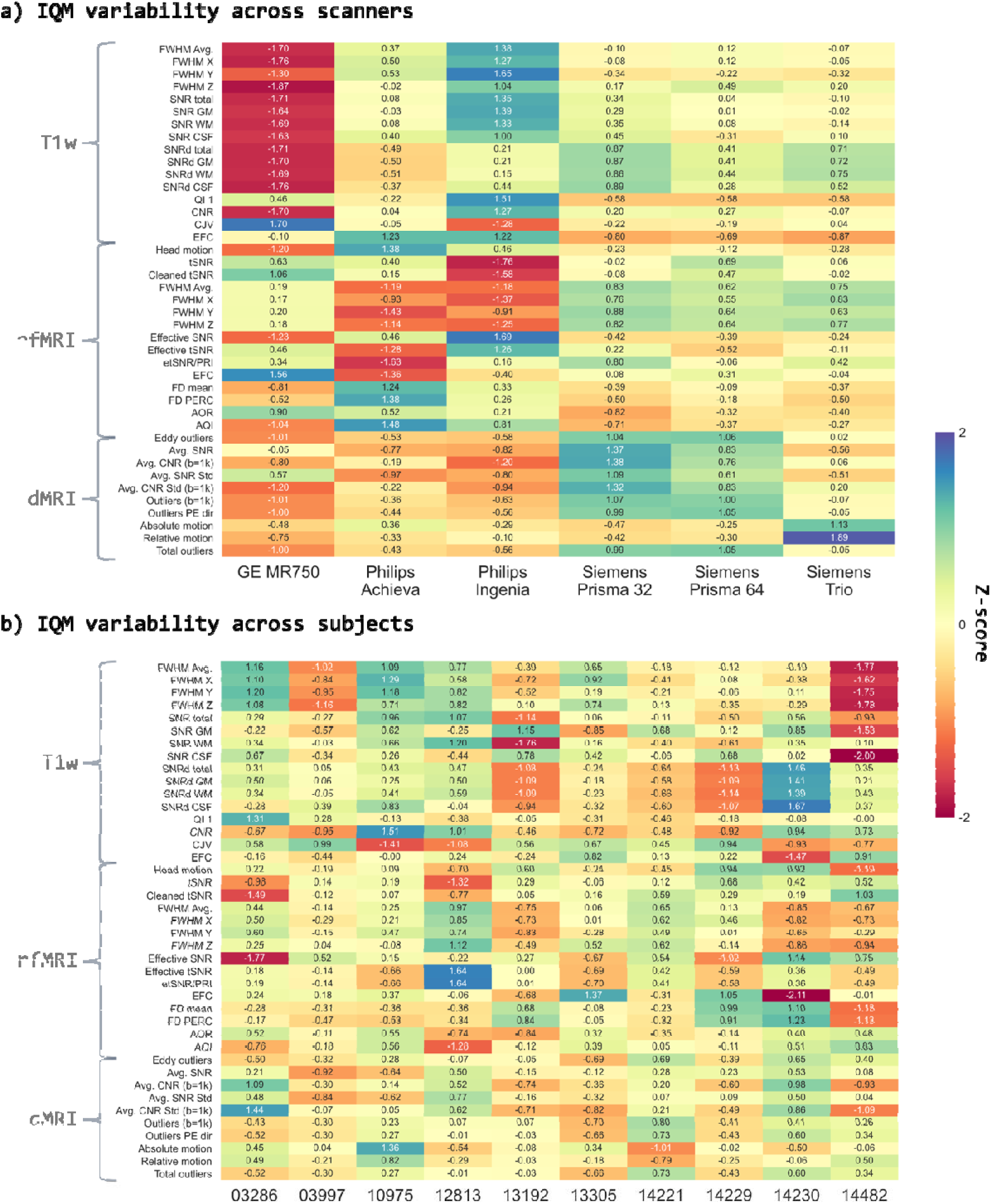
Heatmaps of Image quality metrics (IQM) variability. Top: IQM variability across scanners. Each quality metric for each subject was z-scored across the six scanners. The Z-scores were then averaged across the 10 subjects. Bottom: IQM variability across subjects. Each quality metric was z-scored across subjects and then averaged across the six scanners. In each case, we exclude within-scanner repeats. Higher positive or negative values represent large deviations from the mean z-scored IQM across scanners/subjects. We were unable to acquire multi-shell data for all scanners, hence we exclude higher b-value IQMs in these comparisons.

### 3.2 Mapping Between-Scanner Variability for Multi-Modal IDPs

We subsequently used the data to extract multi-modal IDPs and explore their between-scanner variability. First, we used the IDPs to assess between-session similarity. To do so, we initially looked at individual IDP categories and calculated the Spearman’s rank correlation 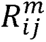 (see Eq. 2.1) for each IDP category between all session pairs (Supplementary Figure 4). Between-session similarity matrices based on T1w-derived IDPs had larger correlation values and tended to be more structured overall, but more so for some IDP categories than others, e.g. within-subject similarity was higher than between-subject for FreeSurfer cortical features, but less so for subcortical ones. This pattern was also present for correlation matrices derived from dMRI IDPs, although the magnitude of correlation values was typically reduced. Correlation matrices derived from fMRI IDPs were less structured and had considerably lower correlation values.

Subsequently, we took the median across IDP categories (Eq. 2.1) to obtain an overall between-session similarity metric considering all IDP categories for each session (Supplementary Figure 5). The pattern previously described was apparent. In addition, we also observed how within-scanner repeats of the same subject were more similar than between-scanner repeats of the same subject, highlighting the harmonisation challenge. To better visualise these differences, we focused on the sessions of the four subjects that had both between and within-scanner repeats (Figure 4, left). This qualitatively demonstrates greater similarity for within-scanner repeats (blue outline) compared to between-scanner repeats (green outline). This is confirmed when comparing the distribution of between-session correlation values (Figure 4, right), illustrating a greater consistency in values of IDPs derived from within-scanner measurements compared to those derived from between-scanner data. Importantly, we also observe an overlap in correlation distributions for between-subject-within-scanner sessions and within-subject-between-scanner repeats. This indicates that IDP similarity for the same subject scanned on different scanners may be as low as the IDP similarity for different subjects scanned on the same scanner.

**Figure 4.**
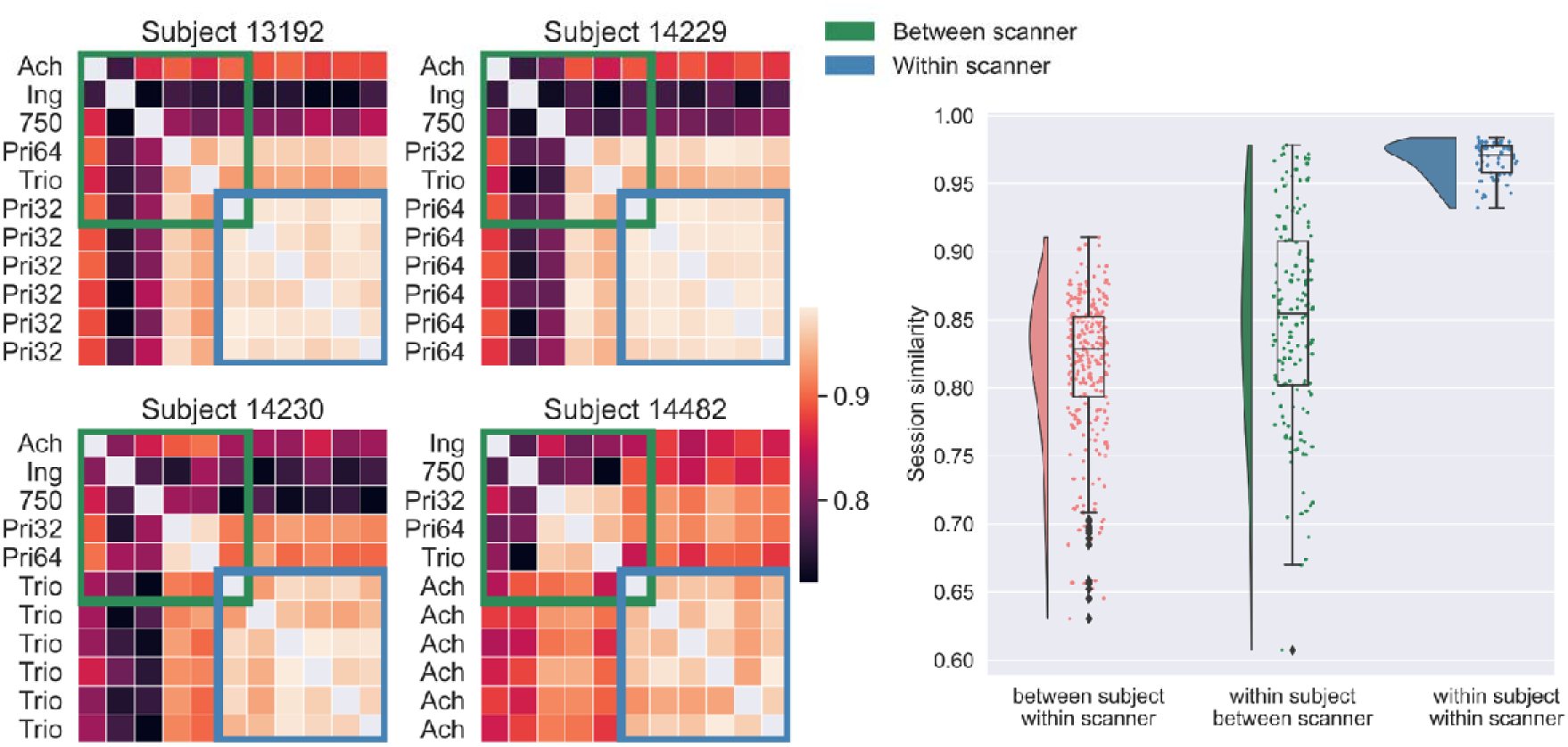
Between-session similarity. Left: Correlation (Spearman’s rank) matrices (see Eq. 2.1) depicting the similarity of IDPs between scanning sessions for the four subjects with within-scanner repeat scans. Spearman’s rank correlation is calculated between all session pairs for IDP categories (Supplementary Figure 4) and the median across categories (Supplementary Figure 5) is presented for the subset of subjects. IDP categories include subcortical volumes, brain tissue volumes, subcortical T2*, cortical parcel volumes, dMRI regional and tract-wise microstructure (FA, MD, MO, L1, L2, L3), rfMRI functional connectivity node amplitude and edges. Right: the distributions of within/between scanner/subject session similarities.Each data point represents the median (across IDP categories) correlation between a pair of sessions, i.e. entries of.

We subsequently explored, for each IDP, the presence of scanner-related bias, by checking how the mean values for that IDP across between-scanner repeats agreed against the mean across within-scanner repeats (Figure 5) ([between-scanner mean – within-scanner mean]/within-scanner mean, expressed as percentage). Even if the differences were larger for some dMRI-extracted IDPs and considerably higher for fMRI-extracted IDPs, bias was consistent and relatively low across the group-level and subject-level and mostly in the range of *±*10%.

**Figure 5.**
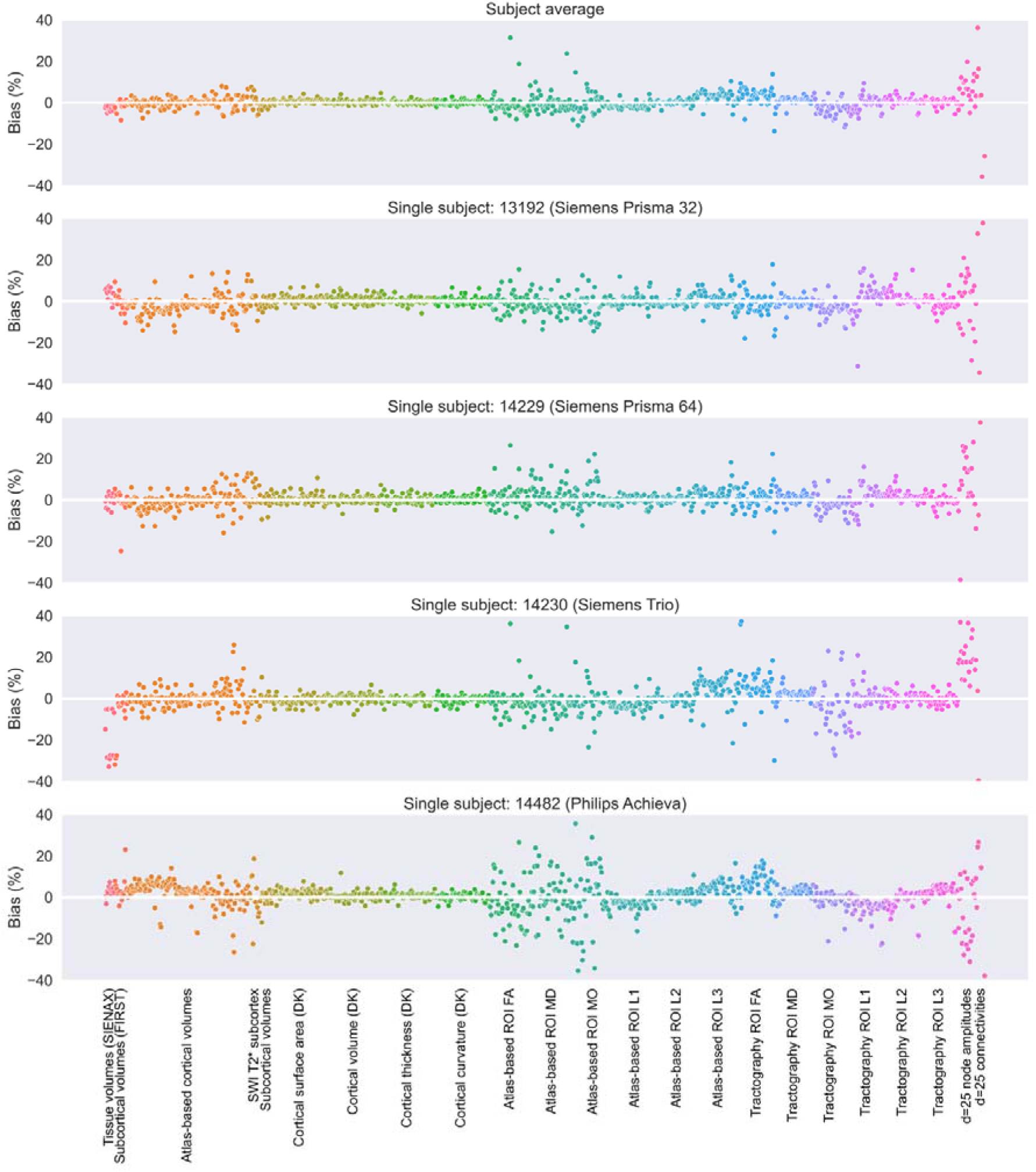
The bias of imaging-derived measures comparing the mean of between-scanner measurements from six different scanners and six within-scanner measurements, reflecting the average across four subjects/scanners (top), the average across scanners from the same vendor (second row), and single subjects (rows 3-6). Bias is calculated IDP-wise as the relative difference between the mean of the between-scanner repeats and the mean of the within-scanner repeats, i.e. 100*(between scanner mean – within scanner mean)/within scanner mean.

We then explored how within-subject between-scanner variability for all considered IDPs compares against two baselines: a) within-scanner variability, b) between-subject (biological) variability. Figure 6 shows the CoVs for each IDP for within-scanner repeats and for between-scanner repeats. Plotted together (third row), and by comparing IDP-group means (fourth row), it becomes apparent that the between-scanner variability can be on average as large as ∼5 times the within-scanner variability, as confirmed by the relative difference (fifth row). We also compared between-scanner repeat variability to “biological” variability (between-subject-within-scanner: orange in rows three and four), and we found that the between-scanner variability is not always smaller than the biological variability (bottom row) for several of the IDP groups. IDP-group-wise medians in the relative difference (rows five and six) are reported in Supplementary Table 5. Certain IDPs (e.g. T1w-extracted atlas-based parcellation IDPs) showed between-scanner variability exceeding 5 times that of the within-scanner variability and over twice that of biological variability. At the IDP-group level, the median between-scanner CoV exceeds a relative difference of 200% in 6 of 23 IDP groups when comparing against within-scanner repeat variability (Figure 6, fourth row). Comparing to biological variability, median between-scanner CoV exceeds that of biological variability in 5 of 23 IDP groups (Figure 6, fifth row).

**Figure 6.**
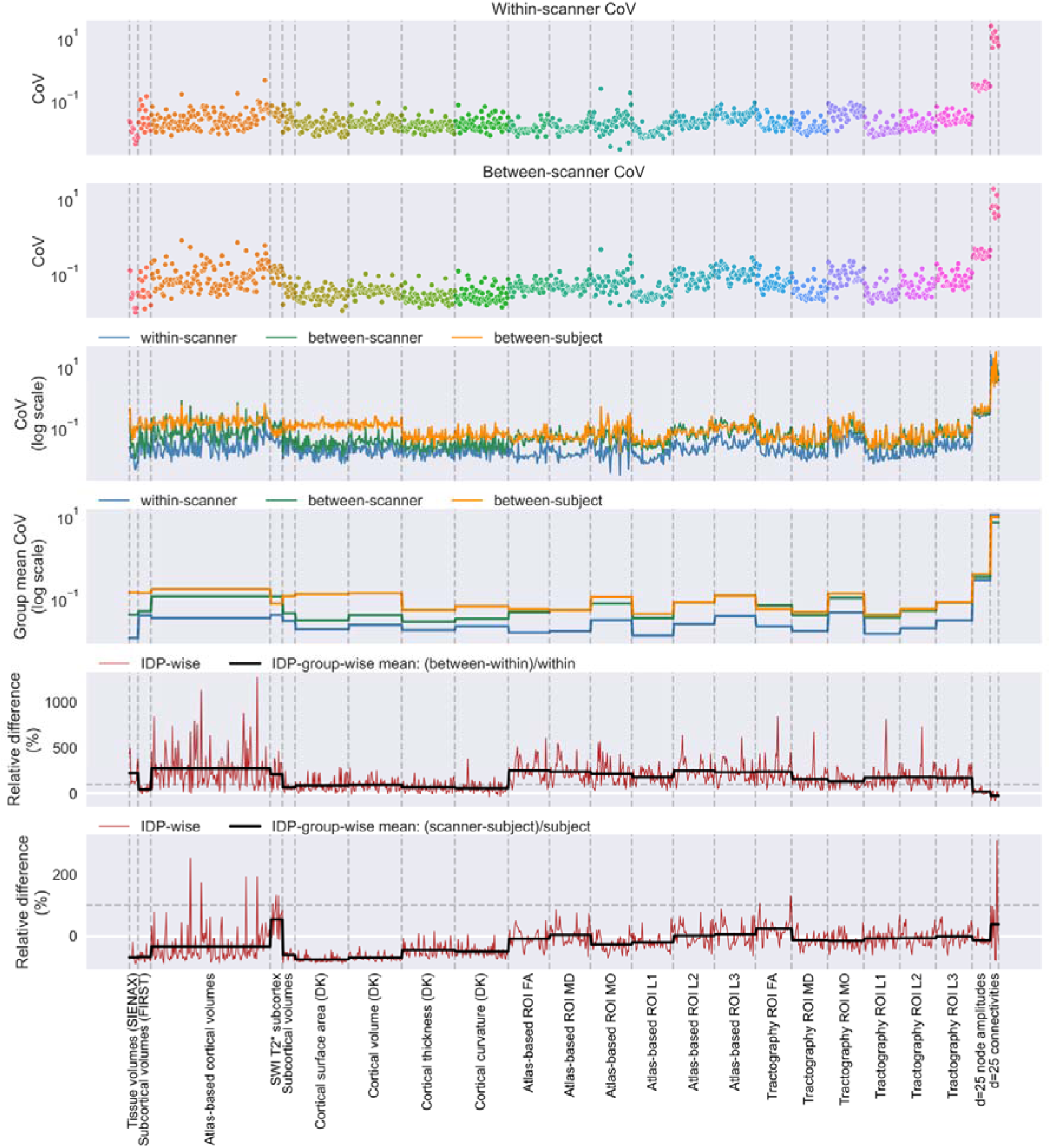
The coefficients of variation (CoVs) of IDPs within/between scanner repeats. Top row: the IDP-wise CoVs across 6 within-scanner repeats, averaged across the four subjects with within-scanner repeats. Second row: the IDP-wise CoVs across six between-scanner repeats, averaged across all subjects. Third row: the within-scanner (blue), between-scanner (green) and between-subject-within-scanner (orange, reflecting biological variability) CoVs plotted on a log-scale. Fourth row: the IDP-group-wise mean of the CoVs (from the third row) plotted on a log scale for within-scanner (blue), between-scanner (green) and between-subject-within-scanner (orange) sessions. Fifth row: the IDP-wise (red) and IDP-group-wise (black) relative difference (between-within/within [scanner]) in CoVs. Bottom row: the IDP-group-wise relative difference in between-scanner CoVs (within scanner, blue; between-scanner, green) and between-subject (biological) CoVs. The dashed horizontal line in rows five and six indicate relative difference of 100%.

We observed trends in variability not only relating to the modality from which the IDPs are derived, but also to the type of processing used to derive said IDPs. For instance, T1w-extracted atlas-based parcellation IDPs show greater between-scanner variability compared to T1w-extract FreeSurfer IDPs, reflecting sources of variability introduced in the processing pipeline. Whilst dMRI-extracted IDPs show relatively high between-scanner variability, they are relatively consistent across processing methods although with reduced variability on average for the tractography-based IDPs compared to the atlas-based IDPs, and with some expected trends. For example, between-scanner variability for both atlas-based and tractography-based IDPs is larger for L3 compared to L2 and compared to L1. IDPs extracted from the NODDI-modelled dMRI data generally have higher between-scanner variability compared to those extracted from the DTI model (Supplementary Figure 7). IDPs derived from SWI showed high between-scanner variability, exceeding biological variability, but a within-scanner variability comparable with other IDP groups. rfMRI-extracted IDPs were particularly variable, with connectivity edges showing very high variability for both biological and scanner related variability and within-scanner variability exceeding biological variability. A version of Figure 6, but using only the four subjects with within-scanner repeats when calculating the between-scanner CoVs, is provided in Supplementary Figure 6, revealing very similar trends.

For each IDP, we also explored the consistency in subject ranking across scanners (Figure 7). A value of 1 indicates perfect consistency, i.e. all ten subjects are ranked in the same way when using the same IDP across the different scanners. As expected, we see that ranking is preserved more for scanners from the same vendor, with it becoming less consistent when we include scanners from different vendors. However, there are only a few categories of IDPs that are close to the ideal consistency described above. Furthermore, the extent to which ranking is preserved depends on the imaging modality. Between-subject ranking is preserved the most for IDPs from anatomical imaging modalities, followed by susceptibility and diffusion, and the least for functional modalities.

**Figure 7.**
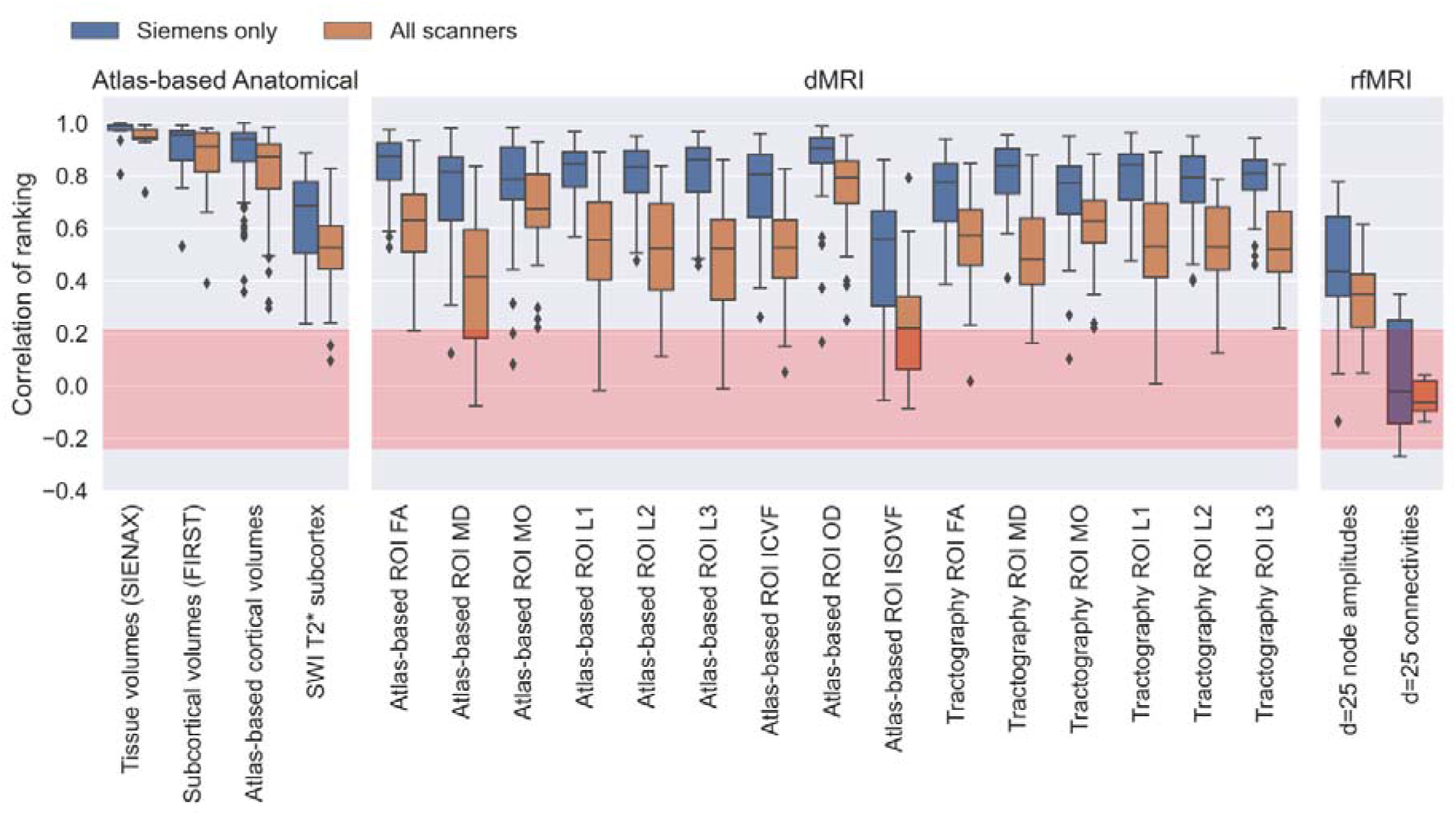
Between-scanner consistency of subject ranking (see Eq. 2.2) for all IDPs grouped by IDP category. The Spearman’s rank correlation is calculated across subjects for each scanner, both for all scanners and restricted to scanners from the same vendor (Siemens). The red region depicts the null distribution’s interquartile range.

To summarise, our database reveals interesting patterns of between-scanner non-biological effects and demonstrates the important need for harmonisation in hundreds of multi-modal IDPs. In the following section, we explore how our database can be used as a testbed for both implicit and explicit harmonisation approaches.

### 3.3 A Testbed for Evaluating Harmonisation Approaches

#### 3.3.1 Implicit Harmonisation

Our data can also be used to assess the robustness of processing pipelines when applied to data from different scanners and compare alternatives for extracting similar IDPs. In this section we demonstrate three examples of such pipeline optimisation, (i) for extracting cortical area volumes from anatomical images, (ii) for extracting subcortical volumes from anatomical images and (iii) on the effect of dMRI denoising on DTI metrics.

We first explored how different approaches for obtaining cortical area volumes (i.e. atlas-based vs FreeSurfer vs FastSurfer) affect between-scanner variability of volumetric IDPs, using the within-scanner variability as a baseline (Figure 8a). To do so, we compared the CoV and consistency of subject ranking for cortical area volumes derived using an atlas-based registration approach (with 96 parcels, as done in the UK Biobank pipeline) to those derived from FreeSurfer (DK coarse with 63 parcels, and Destrieux fine with 148 parcels) and FastSurfer (coarse DK only). We found comparable within-scanner variability across approaches, although with greater variability for the fine FreeSurfer (Destrieux) parcellation scheme. However, between-scanner variability is mostly consistent with the within-scanner variability for the two FreeSurfer-based approaches (DK: 0.033 compared to 0.019; Destrieux: 0.049 compared to 0.037 for median between-scanner and within-scanner respectively), followed by FastSurfer (0.048 between-scanner and 0.019 within-scanner) and lowest for the atlas-based approach (0.056 between-scanner and 0.015 within-scanner). When considering the consistency of subject ranking, a similar trend is observed (though with numbers inverted as the best rank correlation is high, not low), with the atlas-based (median correlation 0.91) and fine FreeSurfer (0.88) parcellation IDPs showing worse ranking consistencies compared to the coarser FreeSurfer/FastSurfer (0.93 and 0.92 respectively) parcellation volumes.

**Figure 8.**
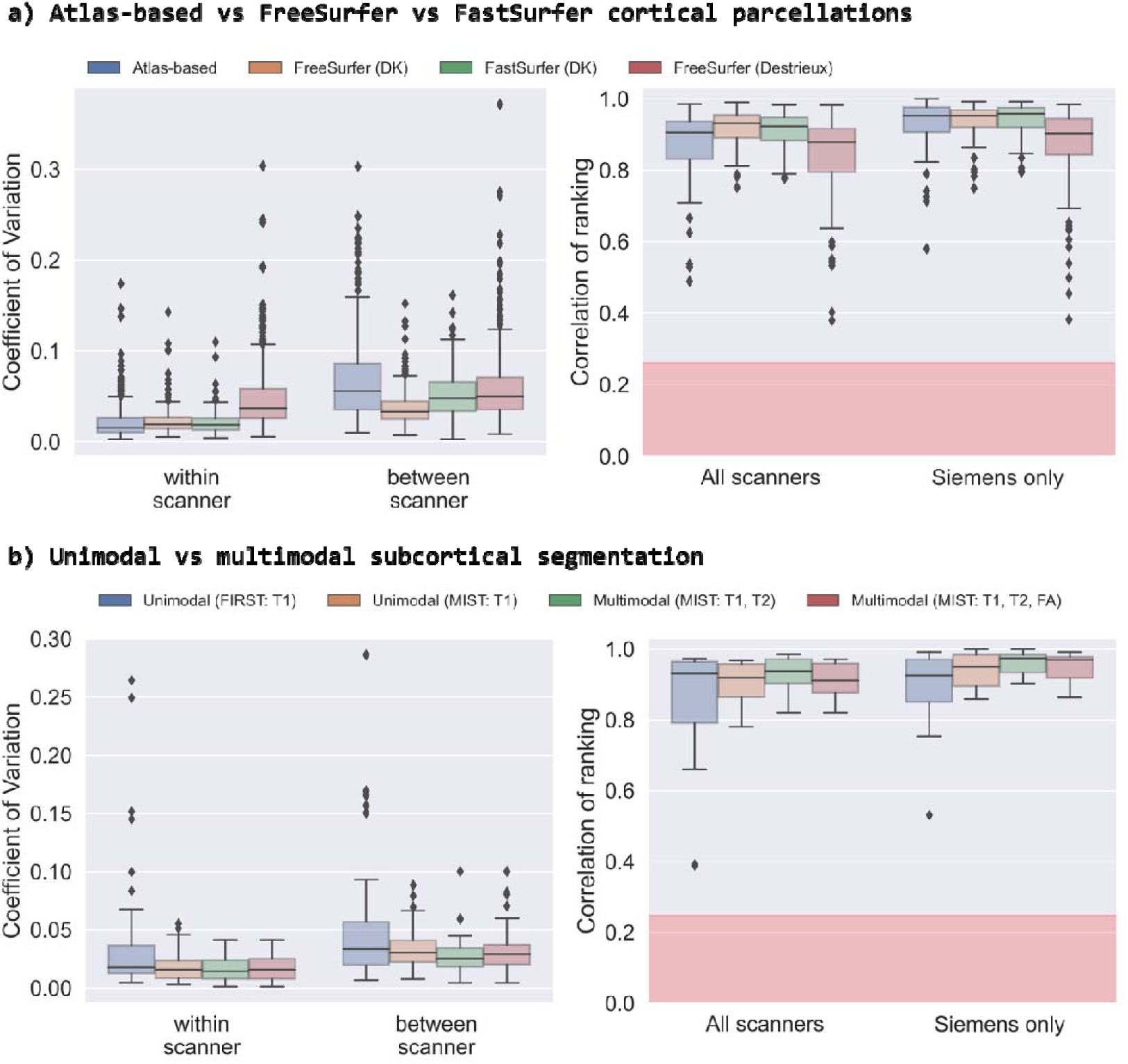
Reproducibility of cortical and subcortical segmentations using different approaches. a) Comparing cortical volumes derived through (i) the registration of an atlas-based parcellation scheme (96 parcels), (ii) FreeSurfer using the Desikan-Killiany parcellation (63 parcels), (iii) FastSurfer with the Desikan-Killiany parcellation, and (iv) FreeSurfer using the Destrieux parcellation scheme (148 parcels). b) Comparing subcortical segmentation volumes derived through (i) unimodal (T1w) segmentation with FIRST, (ii) unimodal (T1w) segmentation with MIST, (iii) multimodal (T1w and T2w) segmentation with MIST, and (iv) multimodal (T1w, T2w and dMRI-derived FA map) segmentation with MIST. In each case, we compare the within-/between-scanner coefficients of variation and the consistency of subject ranking across approaches. The red regions depict the null distribution’s interquartile range.

As a second example, we compared the consistency of ROI-wise subcortical volumes derived using a range of segmentation algorithms, specifically unimodal (using FIRST and single-modality MIST) and multi-modal (using two/three modalities with MIST) segmentation. Figure 8b shows that unimodal segmentation with FIRST and MIST are more variable, for both within-scanner (0.018 and 0.016 respectively) and between-scanner (0.033 and 0.030 respectively) repeats, and show the lowest consistency in subject ranking (median correlation 0.93 and 0.92 respectively). Multimodal subcortical segmentation with MIST (using two anatomical modalities) achieves the best consistency when comparing between-scanner and within-scanner variability (0.025 and 0.015 respectively) and high subject ranking consistency (median correlation 0.94).

We next used our resource in a slightly different way, capitalising on the availability of multiple within-scanner repeats. We explored the effect of denoising on dMRI data, anticipating that since thermal noise is a major contributor to within-scanner variability, denoising the data should lead to a reduction in within-scanner variability of IDPs compared to raw (not denoised) data. When considering tract-wise averaged DTI metrics (FA/MD) across within-subject-within-scanner repeats, Figure 9a demonstrates that denoising induces relatively small differences, most likely reflecting relatively high SNR in the data. Even if, for a number of IDPs, variability was reduced with denoising, this was not always the case, contrary to expectation. We observed IDPs, particularly for tracts in inferior regions (cerebellum, brainstem, uncinate fascicle) where within-scanner variability without denoising was smaller than the one obtained from denoised data. As the type of denoising that we performed is patch-based and the main processing that occurs after denoising and before the extraction of IDPs is distortion correction (including susceptibility-induced distortion corrections), we explored whether these counter-intuitive results in the inferior parts of the brain were related to distortion levels that are higher in these brain regions. We found that regional off-resonance frequency (which is proportional to the amount of distortions) explains some of this behaviour (Figure 9b, moderate correlations that are statistically significant), hinting at interactions between patch-based denoising and distortion correction. We hence re-processed the data and denoised it only after distortion correction. This approach is suboptimal as it changes the statistical properties of the signal and violates assumptions that denoising methods rely on, hence it is not suggested in the general case. Nevertheless, it was used here as a confirmatory test, since it reduces potential interactions between the denoising patches and the shape corrections performed to reverse susceptibility-induced distortions. In doing so, we found reduced associations between the relative difference in CoVs and regional off-resonance frequency (Figure 8c, magnitude of correlations dropped and statistical significance was no longer observed).

**Figure 9.**
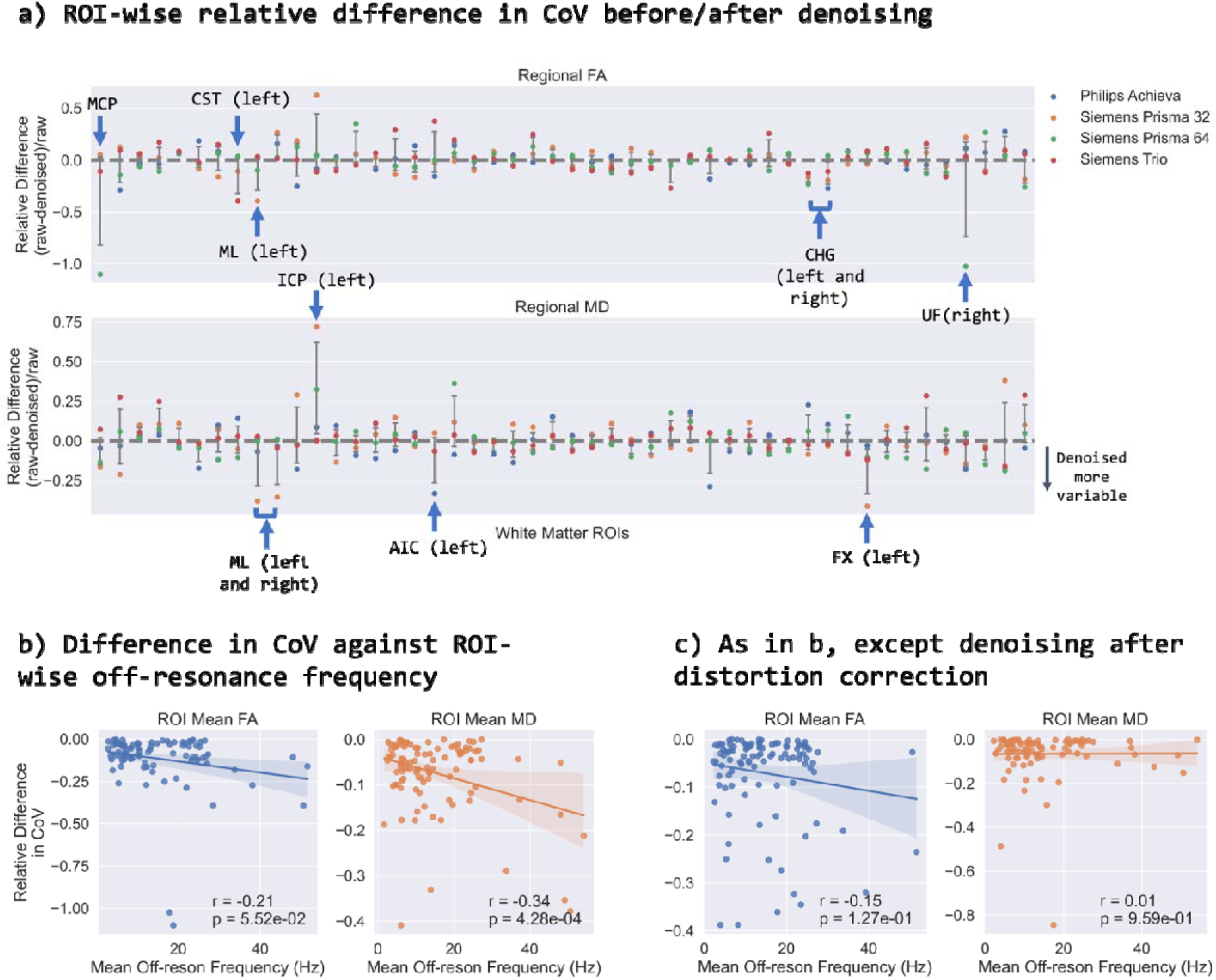
The effect of denoising on tract-wise (TBSS) IDPs. a) The relative difference in region-wise CoV before and after denoising for tract-wise mean FA (top) and MD (bottom). CoV for each IDP is calculated for each subject across the six within-scanner repeats and plotted for each tract. Grey bars represent the mean and standard deviation across the four scanners. b) The session-wise tract-wise CoV against tract-wise mean off-resonance frequency (absolute value in Hz) for regions showing more variability after denoising. c) As in b, except here, we perform denoising after distortion correction.

In summary, these results highlight the importance of carefully considering the different steps in processing pipelines and how data resources like the one presented here can provide important testbeds towards better understanding the implications of processing choices.

#### 3.3.2 Explicit Harmonisation

In addition to the implicit harmonisation examples presented before, we used the data to evaluate existing harmonisation approaches, using the within-scanner variability as a baseline. These approaches are meant to explicitly reduce between-scanner variability. We applied ComBat and CovBat to a number of multi-modal IDPs, including atlas-based cortical area volumes, subcortical volumes obtained from FIRST, ROI-averaged T2* values extracted from susceptibility-weighted images and the FA of white matter ROIs obtained from diffusion MRI. We compared the between-scanner CoV before and after harmonisation (Figure 9, top), with within-scanner CoV as a baseline, and the consistency of subject ranking before and after harmonisation (Figure 10, bottom). In all cases, the CoVs were greater for between-scanner repeats compared to within-scanner repeats. Both harmonisation approaches reduced the between-scanner variability towards the within-scanner variability baseline in each set of IDPs. Success in doing so is variable across IDPs. For instance, ComBat worked better in harmonising SWI T2* values compared to atlas-based cortical area volumes. Interestingly, however, and common across all IDPs, between-scanner subject ranking consistency before and after harmonisation was almost identical. ComBat and CovBat modify IDP values such that variability is reduced but they are not beneficial for improving cross-subject ranking between scanners. This is not the case for the pipeline modifications presented in the previous section, suggesting that blindly performing explicit harmonisation without carefully considering processing pipelines may be suboptimal, and that a combination of explicit and implicit methods is desirable.

**Figure 10.**
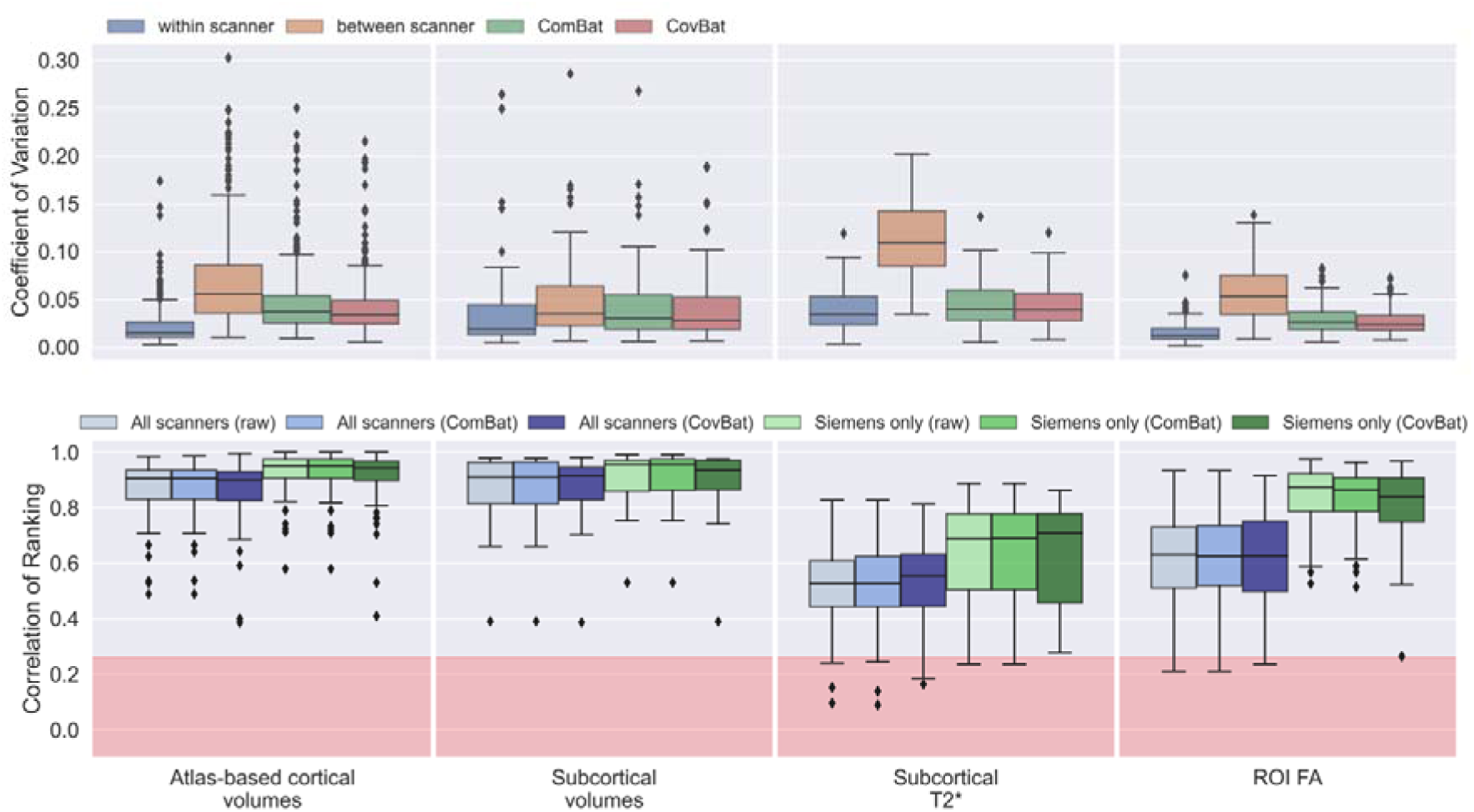
The effect of harmonising IDPs using ComBat and CovBat. Top: the IDP-wise coefficient of variation (CoV) before and after harmonisation. For each of the four subjects with within-scanner repeats, CoV was computed for each IDP across the six repeats (either within-scanner or between-scanner), prior to harmonisation. After harmonisation, the IDP-wise CoV is calculated for the between-scanner repeats. Bottom: the IDP-wise correlation of subject ranking before and after harmonisation for scanners of the same vendor and for all scanners. The red regions depict the null distribution’s interquartile range.

## 4. Discussion

We have presented a comprehensive harmonisation resource for multi-modal neuroimaging data, based on a travelling-heads paradigm. We have used this to map between-scanner effects across hundreds of multi-modal IDPs and shown that between-scanner variability is up to 10 times larger than within-scanner variability of the same modality IDPs for the same subject. Importantly, for a number of IDPs, between-scanner variability can be of the same size as between-subject (“biological”) variability. We also found that consistency in subject ranking across scanners can be compromised relatively easily, particularly for certain modalities and IDPs. Compared to previous travelling-head studies (Pohl et al., 2016; Tax et al., 2019; Yamashita et al., 2019; Tong et al., 2020; Maikusa et al., 2021; Kurokawa et al., 2021; Tanaka et al., 2021; Duff et al., 2022; Tian et al., 2022), our study provides a more comprehensive harmonisation resource in a number of ways: (i) data is acquired from all 3 major vendors and from different generations of scanners from the same vendor, (ii) data is acquired from different imaging sites where radiographers and practices are different, (iii) data is acquired from many neuroimaging modalities, (iv) multiple scan-rescan data is acquired which allows the assessment of within-scanner, within-subject variability in addition to between-scanner variability, and (v) hundreds of multi-modal IDPs are considered using a modified and augmented version of the UK Biobank pipeline. Our resource is publicly released (made available on publication) and will be augmented with further subjects and scanners in the coming years, including additional within-scanner repeats (two new GE MR Premier wide-bore scanners are already installed in two different sites of our study).

Our resource has been designed to allow for different baselines to compare between-scanner effects: multiple within-scanner-within-subject repeats to capture within-scanner variability baselines and multiple subjects to capture between-subject (biological) variability. We found that IDPs derived from T1w imaging are, in general, the most consistent, but we also observed that this heavily depends on the processing approach. These were followed by IDPs derived from dMRI yet, even within these IDPs, there was a spectrum of variabilities depending on the type of measure (e.g. NODDI more variable than DTI, atlas-based more variable than subject-specific tractography). The IDPs derived from rfMRI were most variable. These trends are consistent with findings of other recent multi-modal studies that considered fewer scanners (Duff et al., 2022). We have also shown that the least between-scanner variability is observed when using scanners from the same vendor, as anticipated. Introducing different vendors increases the variability in IDPs and also decreases consistency in ranking of subjects across scanners.

Previous work has reported similar trends to the ones reported here. For instance, structural IDPs were the most reproducible of the IDPs we present, and this is consistent with past findings. High repeatability of these IDPs has been shown across a range of segmentation approaches (de Boer et al., 2010), across multiple sites (Jovicich et al., 2006) and across scanners of varying magnetic field strength (Fujimoto et al., 2014). Cortical areas and volumes derived from FreeSurfer have been shown to even be robust to different acquisition sequences (Knussmann et al., 2022). It is worth noting that among the various groups of structural IDPs, a previous study (Duff et al., 2022) has shown that cortical area and thickness as derived from FreeSurfer are more robust than the grey matter volumes which were estimated for 139 ROIs and this is in agreement with our findings.

For diffusion related IDPs, previous studies have shown that generally, NODDI parameters have larger between-subject variations than DTI IDPs (Chung et al., 2016, p. 216; De Luca et al., 2022). The CoV for ISOVF has been observed to be consistently the largest among diffusion IDPs (Chung et al., 2016, p. 216), which is in agreement with our results (Supplementary Figure 7). Of the DTI IDPs, FA has been found to be less robust than MD (Chung et al., 2016, p. 216; Farrell et al., 2007), as it reflects a higher moment of the tensor eigenvalues. This in agreement with our results, which also show that L1, which is larger in magnitude, is less sensitive to between-scanner effects that the smaller L2 and L3. Methods have recently been developed specifically to harmonise IDPs derived from higher-order dMRI models (De Luca et al., 2022).

For rfMRI IDPs, it has been reported previously that test-retest reproducibility is a limiting factor (Castellanos et al., 2013), which also explains the large relative variability values we found. The results we have presented demonstrate that the difference in the variability of between-vs within-scanner repeats in rfMRI was low, since within-scanner variability was already high. Other studies that performed similar analyses (Duff et al., 2022) pointed out that IDPs reflecting pairwise connectivity (as well as node amplitudes) do not show a high level of reliability across sites, therefore consistency in summary ICA components was instead evaluated. Furthermore, in the study performed by (Jovicich et al., 2006), significant inter-site differences in connectivity scores were found.

We demonstrated how our resource can be used as a testbed to explore and evaluate harmonisation approaches. The existence of multiple within-scanner repeats allowed us to define a consistent and interpretable reference to compare harmonisation efficacy against and avoided the need to use ad-hoc methods, such as group matching based on covariates (Fortin et al., 2018; Garcia-Dias et al., 2020), for validation. Specifically, we have shown how our data can be used to optimise processing steps used in IDP extraction pipelines (implicit harmonisation), such that between-scanner variability in extracted IDPs is minimised compared to e.g. within-scanner variability. We have also tested the performance of post-processing harmonisation tools (explicit harmonisation) and specifically checked whether the harmonised IDPs are indeed less variable between-scanners (and by how much) compared to no harmonisation. Overall, we found that even though the tested explicit harmonisation methods tested did remove parts of non-biological variability, they did not recover inconsistent cross-subject ranking across scanners. This was not the case for implicit harmonisation methods, suggesting that a consideration of both is needed to achieve optimal results.

More specifically, for anatomical IDPs, we found that cortical area volumes extracted from FreeSurfer and subcortical volumes extracted from multi-modal segmentation have between-scanner variability that is closer to the respective within-scanner variability (and hence are less sensitive to between-scanner effects) compared to other approaches explored. Previous studies have shown that cortical volumes derived from FreeSurfer have a strong degree of robustness against scanner effects. For instance, in (Iscan et al., 2015) it is shown that for the DK atlas, cortical volume measures showed test-retest correlation scores (from scans acquired at four different sites) of 0.88. This study also showed higher test-retest correlation and inter-class correlation scores for volumes from the DK atlas (coarse) than the Destrieux atlas (fine), which is in agreement with the results we obtained. These results confirm what we expect since regions defined by the DK atlas are larger than those in the Destrieux atlas.

For subcortical volumes, we found volumes derived using a multi-modal segmentation method (MIST) were more reproducible than those derived using a unimodal approach (FIRST). This is in agreement with the findings in (Visser et al., 2016) who compared the approach with FIRST and FreeSurfer using a manual segmentation as a benchmark. We also assessed the advantage of using MIST with data from three modalities (T1w, T2w and dMRI data) compared to training it using two modalities (T1w and T2w), and in a unimodal fashion (T1w only). Intuition would suggest that leveraging imaging information from more modalities would result in more reproducible results, however, our results show that adding dMRI data as an input to MIST decreased between-scanner reproducibility. These findings agree with results in (Visser et al., 2016), who found that increasing the number of modalities used for MIST segmentation can increase variability. This can happen for regions where the contrast is very clear from structural images. In this case, segmentations from the structural images alone are highly reproducible and adding another modality, particularly a more noisy one like dMRI, introduces new sources of variability.

We found a slightly unexpected trend for dMRI denoising using MP-PCA (Veraart et al., 2016). Within-scanner variability of extracted dMRI IDPs did not always decrease after denoising compared to IDPs extracted from “raw” data. It is worth pointing out that raw SNR and CNR values do increase after denoising in this data (Supplementary Figure 8). The natural question to ask is why then does the variability of these IDPs does not improve after denoising? A possible explanation is that we observed highly variable IDPs in the caudal regions of the brain where denoising appeared to have increased the variability. These are areas known to be prone to susceptibility artefacts (Andersson et al., 2003) and therefore distortion correction is more impactful in these areas. The fact that we see these areas significantly affected after denoising suggests that there is a possible interaction between denoising and distortion correction (Figure 9b). This could happen because, even prior to distortion correction, denoising assumes that every voxel is in the correct place yet this is not true in the presence of distortions. As denoising is patch-based, incorrectly placed voxels would end up influencing the denoising process meaning a distortion correction like this could lead to misplaced voxels and in slightly different ways for the different repeats. To further explore this, we applied denoising after distortion correction and found a reduced association between differences in variability and off-resonance frequency (Figure 8c). However, we should note that by applying distortion correction prior to denoising will break some of the assumptions in the MP-PCA algorithm. These findings suggest that the optimal way of denoising requires more exploration and suggests that denoising and distortion correction may ideally have to be considered simultaneously (similar in spirit to the simultaneous consideration of all distortion fields and their correction in (Andersson and Sotiropoulos, 2016)).

We also compared explicit harmonisation approaches in ways that have not been evaluated before. We showed that both ComBat (Fortin et al., 2017) and CovBat (Chen et al., 2022) reduced the between-scanner variability for a range of IDPs derived from different modalities towards the level of the respective within-scanner variability. The relatively small difference in subcortical volumes corrected with ComBat compared to the uncorrected volumes is in agreement with findings from other studies (Treit et al., 2022). The authors in this study used ComBat to reduce systematic variations in the brain volumes of 23 travelling subjects scanned in 3 different scanners and they found minimal changes (of less than 5%) between corrected and raw volumes for several subcortical regions (caudate, globus pallidus, putamen, and thalamus). The authors in (Treit et al., 2022) point out that the degree to which ComBat decreases inter-subject variability likely depends on the magnitude of site effects in the raw data implying that ComBat has less of an effect on results that are more robust to site effects. Our findings support this notion as of the three IDPs tested (subcortical volumes, T2* values and FA values), the subcortical volumes had on average the least between-scanner variability of the three and were also affected the least by ComBat. It is important to note that with 10 subjects and six scanning sessions, we were at the lower end of the recommended sample size for ComBat for independent subjects across different scanners (Fortin et al., 2017), however we are above the minimum suggested requirements for the case of travelling heads (Maikusa et al., 2021).

In summary, we have presented a comprehensive harmonisation resource that we publicly release and will continue to extend in the future. Capitalising on a travelling-heads paradigm and the availability of scanners from all three major MR vendors, the data allow assessment of within/between-subject and within/between-scanner effects. As we have shown, this enables novel evaluations of efficacy of both implicit and explicit harmonisation methods. The resource can be used as a testbed for existing harmonisation approaches, as well as for new ones to be developed in the future.

## Acknowledgements

S.W., A.T. and S.N.S. are supported by an ERC Consolidator grant (101000969 to S.N.S.). A.N. has been supported by the Engineering and Physical Sciences Research Council (EPSRC) and Medical Research Council (MRC) (ONBI CDT). Scan time costs were provided in part by the Nottingham Biomedical Research Centre, by the SPMIC-School of Medicine PhD student and scan time allocation fund and by the WIN Centre, which is supported by a Wellcome Trust Center grant (203139/Z/16/Z). Scanners were operated by local radiographers and physicists (Mr Jon Campbell, Mr Michael Sanders, Mrs Juliet Semple, Mr David Parker, Mrs Caroline Young and Mrs Nicky Aikin for FMRIB Primsa, OCMR Trio and FMRIB OHBA, Mr Andrew Cooper for SPMIC-QMC, Dr Olivier Mougin and Prof Paul Morgan for SPMIC-UP Philips Ingenia and SPMIC-UP Philips Achieva).

## Data and materials availability statement

Anonymised BIDS format data are freely available on OpenNeuro (made available on publication). The adapted UKBB pipeline used is available via GitHub (made available on publication). Jupyter Notebooks used for analyses and summary data are available on GitHub (made available on publication). Software used are freely available.

## Notes

### Competing Interest Statement

The authors have declared no competing interest.

