## Supplementary Information for "A resource for development and comparison of multi-modal brain 3T MRI harmonisation approaches"

#### Total readout time equations

Here, we report the equations used by dcm2niix to calculate the total readout time for each vendor.

**Siemens:** https://github.com/rordenlab/dcm2niix/issues/130

$$EffectiveEchoSpacing = \frac{1}{BWPPPE \times MatrixSizePE}$$

$$TotalReadoutTime = EffectiveEchoSpacing \times (MatrixSizePE - 1)$$

where $BWPPPE$ is the bandwidth per pixel in the phase encoding direction and $MatrixSizePE$ is the dimension of the image reconstruction matrix in the phase encoding direction.

**GE:** https://github.com/rordenlab/dcm2niix/tree/master/GE

$$TotalReadoutTime = \left(ceil\left(\frac{1}{RF} \times \frac{MatrixSizePE}{AssetR}\right) * RF\right) - 1) \times ES \times 0.000001$$

where $RF$ is a round factor applied conditionally such that $RF$ is 2 (Full Fourier) or 4 (Partial Fourier), $MatrixSizePE$ is the image acquisition matrix dimension in the phase encoding direction, $AssetR$ is the in-plane acceleration factor.

**Philips:** https://osf.io/xvguw/wiki/home/

$$ActualEchoSpacing = \frac{WaterFatShift}{ImagingFrequency \times 3.4 \times (EPIFactor + 1)}$$

$$TotalReadoutTime = ActualEchoSpacing \times EPIFactor$$

$$EffectiveEchoSpacing = \frac{TotalReadoutTime}{MatrixSizePE - 1}$$

where $ActualEchoSpacing$ is the acquisition echo spacing, $EPIFactor$ is the equivalent to the echo train length plus 1, $MatrixSizePE$ is the dimension of the image reconstruction matrix in the phase encoding direction.

Note: in our data, the acquisition and reconstruction matrix dimensions are equal.

Minor subject protocol deviations

The majority of subjects were acquired using the same protocols. However, for some subjects there were minor deviations in some protocol parameters that we describe here for completeness.

**GE MR 750 rfMRI:** For one subject (03286), we acquired two versions of the rfMRI data with 1) an isotropic spatial resolution of 2.4mm and 2) a spatial resolution of 2.2x2.2x3.4mm. Neither matched the 3.3mm isotropic data acquired for other subjects. As such, we excluded this subject's rfMRI data from the analyses. For 6 subjects (03286, 03997, 10975, 12813, 14482, 14221) there was a mismatch in PE direction between dMRI and fMRI that we accounted for in the pipeline.

**Philips Achieva dMRI:** In most cases, the dMRI protocol included 6 b=0 s/mm$^2$ volumes. Four subjects (13305, 13192, 14229, 14230) were acquired with 2 b=0 s/mm$^2$ volumes.

**Philips Ingenia dMRI:** In most cases, dMRI data were acquired using an in-plane acceleration factor of 1.5 (TE=98ms, TR=4.4s). For 4 subjects (13305, 13192, 14229, 14230), the dMRI data were acquired using an in-plane acceleration factor of 2 (TE=92ms, TR=3.9s).

Scanner software details

Software version was maintained for the majority of sessions. Siemens Prisma scanners used software version Syngo MR E11 and the Siemens Trio scanner used Syngo B17. The GE M750 used software version DV 24.0. For the Philips scanners, the software version was upgraded during the study. For the Achieva, 6 subjects (03286, 03977, 13192, 13305, 14229, 14230) were acquired with version 5.3.0.3 and the remaining 4 subjects (10975, 12813, 14221, 14482) were acquired with version 5.6.1.1. For the Ingenia, 6 subjects (03286, 03977, 13192, 13305, 14229, 14230) were acquired with version 5.3.1.0 and the remaining 4 subjects (10975, 12813, 14221, 14482) were acquired with version 5.6.1.0.

<u>Supplementary Tables</u>

| **Subject ID** | **Sex** | **Age** | **Within-scan repeats** | **Between-scan interval (days)** | **Within-scan interval (days)** |
|---|---|---|---|---|---|
| 03286 | M | 48 | No | 492 | N/A |
| 03997 | M | 37 | No | 266 | N/A |
| 10975 | M | 25 | No | 455 | N/A |
| 12813 | F | 24 | No | 562 | N/A |
| 13192 | M | 47 | Yes | 314 | 38 (Prisma FMRIB) |
| 13305 | M | 42 | No | 671 | N/A |
| 14221 | M | 25 | No | 555 | N/A |
| 14229 | M | 35 | Yes | 298 | 82 (Prisma WH) |
| 14230 | F | 25 | Yes | 203 | 92 (Trio) |
| 14482 | M | 24 | Yes | 500 | 388 (Achieva) |
| Summary: | 80% M | Mean = 33.2<br>Std = 9.9 | Total = 4 | Mean = 432<br>Std = 153 | Mean = 150<br>Std = 160 |

***Supplementary Table 1** – Subject demographics and time taken to acquire all scans.*

| dMRI | ES (ms) | MB | PI | PF | IF (MHz) | WFS (pix) | BW (Hz/px) | Matrix size PE | EES (ms) | TRT (ms) |
|---|---|---|---|---|---|---|---|---|---|---|
| **Philips Achieva** | 0.67 | None | 2 | 80% | 127.8 | 17.0 | 25.5 | 112 | 0.35 | 39 |
| **Philips Ingenia** | 0.67 | 3 | 1.5 | 80% | 127.8 | 32.7 | 18.1 | 112 | 0.67 | 74 |
| **GE MR750** | 0.68 | None | 2 | None | 127.8 | - | - | 104 | 0.34 | 35 |
| **Siemens Prisma 32ch** | 0.67 | 3 | None | 75% | 123.2 | - | 14.4 | 104 | 0.67 | 69 |
| **Siemens Prisma 64ch** | 0.67 | 3 | None | 75% | 123.2 | - | 14.4 | 104 | 0.64 | 69 |
| **Siemens Trio** | 0.69 | 3 | None | 75% | 123.2 | - | 13.9 | 104 | 0.69 | 71 |

***Supplementary Table 2*** *– The effective echo spacing and total readout times used in distortion correction processing for dMRI data. Values displayed are as reported from dcm2niix (v1.0.20211006) and as used in the equations presented in the Supplementary Information. Where available we report all relevant parameters across vendors. Unavailable parameters are denoted by '-'. We report both the water fat shift and bandwidth for Philips: water fat shift is used in dcm2niix to calculate the effective echo spacing and we report bandwidth to allow comparison across scanners. ES = echo spacing; MB = multiband factor; PI = in-plane acceleration factor; PF = partial Fourier; IF = imaging frequency; WFS = water fat shift (pixels); BW = phase encoding bandwidth per pixel (Hz/pixel); Matrix size PE = image (acquisition and reconstruction) matrix size in the phase encoding direction; EES = effective echo spacing; TRT = total readout time.*

| rfMRI | ES (ms) | MB | PI | PF | IF (MHz) | WFS (pix) | BW (Hz/px) | Matrix size PE | EES (ms) | TRT (ms) |
|---|---|---|---|---|---|---|---|---|---|---|
| **Philips Achieva** | 0.61 | 4 | 1 | 80% | 127.8 | 25.4 | 17.1 | 96 | 0.61 | 58 |
| **Philips Ingenia** | 0.62 | 4 | 1.5 | None | 127.8 | 17.1 | 25.5 | 96 | 0.41 | 39 |
| **GE MR750** | 0.54 | None | 2 | None | 127.8 | - | - | 64 | 0.21 | 13 |
| **Siemens Prisma 32ch** | 0.64 | 8 | None | None | 123.2 | - | 17.8 | 88 | 0.64 | 56 |
| **Siemens Prisma 64ch** | 0.64 | 8 | None | None | 123.2 | - | 17.8 | 88 | 0.64 | 56 |
| **Siemens Trio** | 0.69 | 8 | None | None | 123.2 | - | 16.5 | 88 | 0.69 | 60 |

***Supplementary Table 3*** *– As in Supplementary Table 2 but for rfMRI data.*

| QC Metric | Category | T1w | dMRI | rfMRI |
|---|---|---|---|---|
| Spatial Contrast to Noise Ratio (CNR) | Noise related | / | | |
| Angular Contrast to Noise Ratio (CNR) | | | / | |
| Signal to Noise Ratio (SNR) | | / | / | |
| Temporal SNR (tSNR) | | | | / |
| Coefficient of Joint Variation (CJV) | Outlier and artefact related | / | | |
| AFNI's Outlier index (AQI) | | | | / |
| AFNI's Outlier Ratio (AOR) | | | | / |
| Outliers (Intensity dropout) | | | / | |
| Quality Index 1 (QI 1) | | / | | |
| Absolute Motion | Motion related | | / | / |
| Relative Motion | | | / | |
| Framewise Displacement (FD) | | | | / |
| Eddy Current Distortions | Distortion related | | / | |
| Susceptibility Induced Distortions | | | / | |
| Full-width-half-maximum (FWHM) | Blurring related | / | | / |
| Entropy Focused Criterion (EFC) | | / | | |

***Supplementary Table 4*** *– The image quality metrics (IQMs) used to assess anatomical (T1w), diffusion MRI and functional MRI data quality. IQMs are derived using MRIQC for T1w and fMRI data and eddyQC for dMRI data.*

| IDP group | | Between-scanner vs. within-scanner | | | Between-scanner vs. biological | | |
|---|---|---|---|---|---|---|---|
| | | **Median** | **Quantile (5th)** | **Quantile (95th)** | **Median** | **Quantile (5th)** | **Quantile (95th)** |
| **T1w** | Atlas-based cortical volumes | 223.2 | 34.8 | 731.2 | -55.1 | -78.7 | 60.7 |
| | Subcortical volumes (FIRST) | 31.8 | 3.3 | 112.7 | -69.4 | -82.9 | -43.6 |
| | Tissue volumes (SIENAX) | 134.8 | 104.6 | 464.8 | -71.4 | -89.4 | -42.1 |
| **SWI** | T2* subcortex | 189.9 | 33.6 | 412.8 | 70.0 | -11.1 | 132.1 |
| **T1w FreeSurfer** | Subcortical volumes | 51.3 | 20.2 | 144.3 | -58.0 | -83.4 | -47.8 |
| | Cortical curvature (DK) | 41.6 | -4.6 | 144.0 | -53.4 | -71.7 | -21.1 |
| | Cortical surface area (DK) | 61.2 | 8.3 | 239.2 | -81.0 | -86.0 | -53.0 |
| | Cortical thickness (DK) | 51.3 | -0.9 | 173.7 | -48.3 | -70.7 | 1.5 |
| | Cortical volume (DK) | 80.8 | 9.9 | 205.0 | -72.7 | -81.7 | -50.3 |
| **Atlas-based dMRI** | FA | 212.1 | 100.3 | 472.5 | -11.9 | -52.0 | 38.0 |
| | L1 | 177.6 | 68.3 | 299.9 | -22.2 | -49.2 | 13.3 |
| | L2 | 224.2 | 102.9 | 434.9 | -7.1 | -39.8 | 57.6 |
| | L3 | 219.5 | 89.9 | 396.7 | 0.8 | -45.3 | 64.7 |
| | MD | 215.1 | 101.8 | 472.9 | 0.2 | -32.5 | 70.1 |
| | MO | 180.1 | 57.9 | 453.2 | -30.4 | -63.9 | 16.7 |
| **Tractography-based dMRI** | FA | 216.1 | 80.5 | 442.6 | 21.9 | -34.5 | 83.5 |
| | L1 | 133.7 | 48.0 | 336.8 | -11.6 | -46.8 | 41.1 |
| | L2 | 153.8 | 48.8 | 341.6 | -10.9 | -39.5 | 42.8 |
| | L3 | 174.9 | 53.6 | 321.5 | -2.0 | -41.2 | 48.2 |
| | MD | 134.3 | 40.2 | 282.9 | -16.5 | -48.7 | 16.8 |
| | MO | 129.0 | 44.2 | 226.3 | -18.7 | -50.4 | 32.7 |
| **rfMRI** | d=25 connectivities | -38.1 | -79.4 | 27.8 | 4.4 | -79.2 | 215.8 |
| | d=25 node amplitudes | 10.9 | -9.9 | 61.0 | -18.6 | -27.5 | -1.1 |

***Supplementary Table 5*** *– The mean relative difference in IDP-group-wise coefficient of variation of between-scanner repeats relative to within-scanner repeat and biological variability. Between-scanner vs within-scanner calculated as (between - within)/within. Between-scanner vs biological calculated as (between - biological)/biological.*

### Supplementary Figures

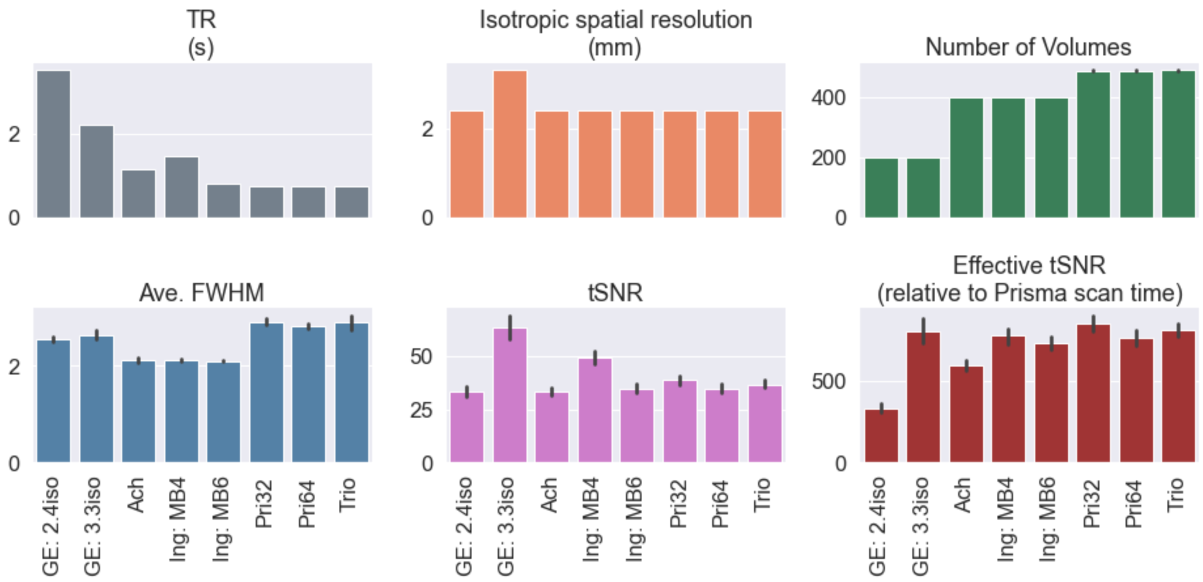

***Supplementary Figure 1*** *– Comparisons of image-quality metrics (IQMs) for different version of the rfMRI protocols across scanners. Each bar represents the mean (and standard deviation – error bars) of the reported IQM across subjects.*

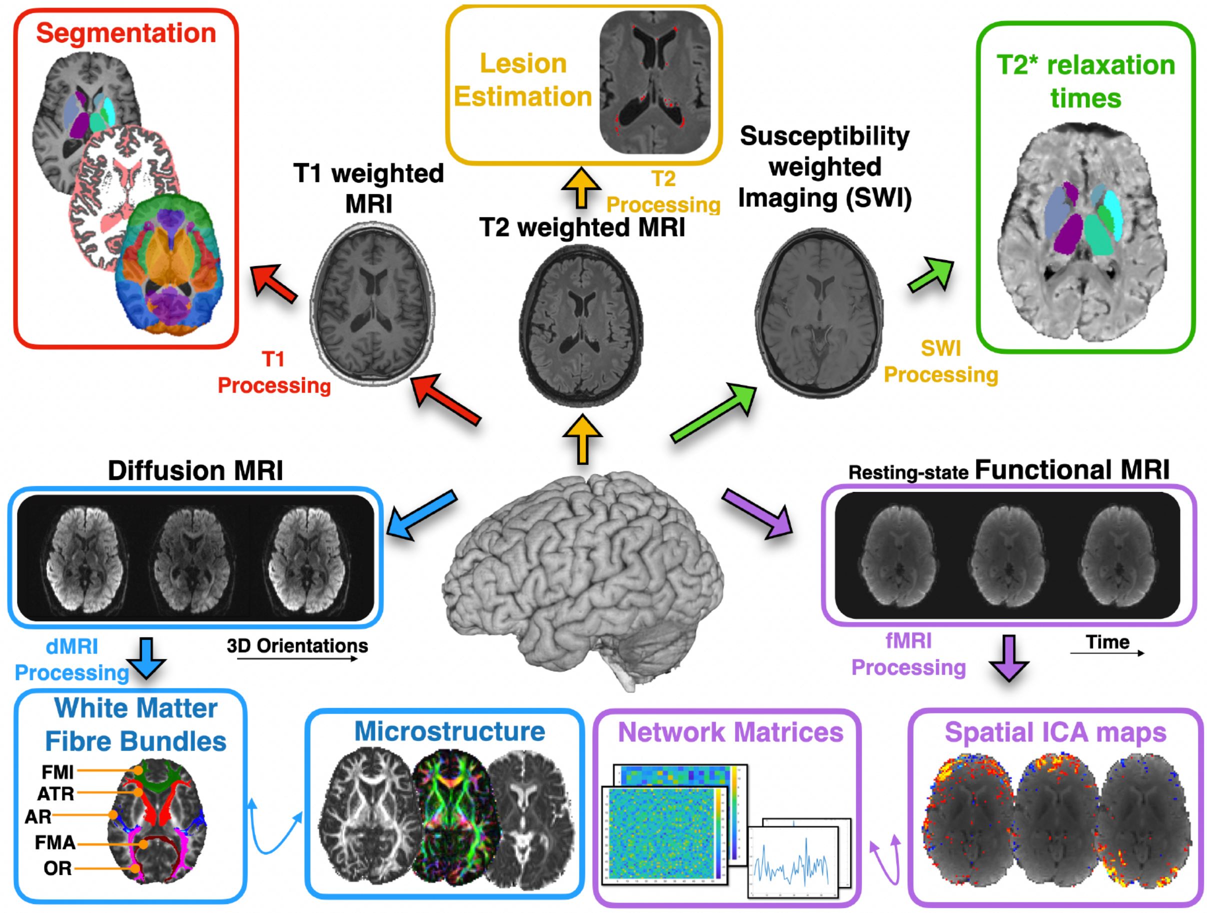

***Supplementary Figure 2 –*** *Overview of the features extracted from each modality. The data from each modality were processed using a modified version of the UK Biobank pipeline to obtain a comprehensive set of imaging features across all scanning sessions.*

A) Variability in fractional anisotropy maps for a single subject within/between-scanners

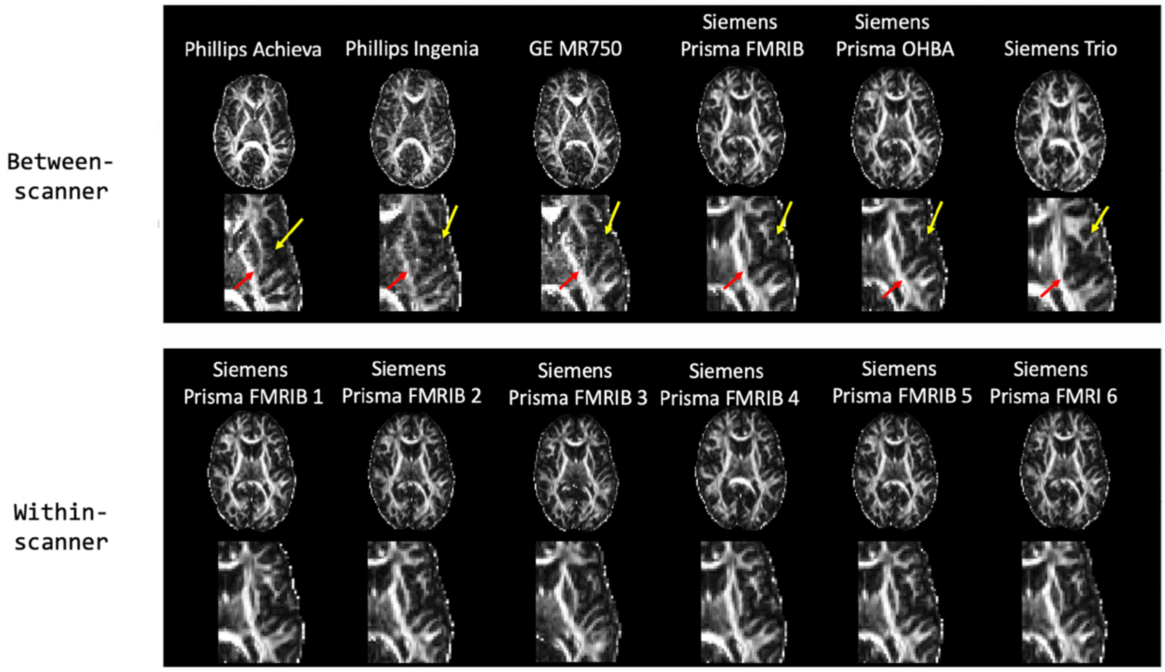

B) Variability in T1w scans for a single subject within/between-scanners

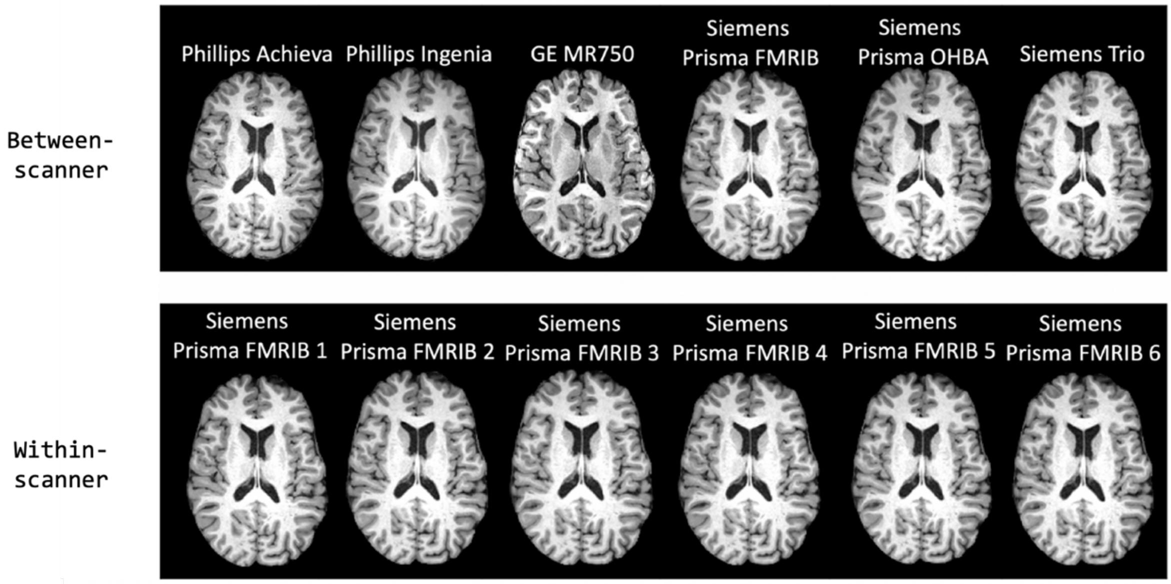

***Supplementary Figure 3*** – *Comparisons of modality-specific variability in quality within/between-scanners. A) Variability in fractional anisotropy (FA) maps between (top) and within (bottom) scanners. Qualitatively, between-scanner variability is greater than within-scanner variability. Specifically, grey/white matter contrast varies appreciably between scanners. Yellow arrows highlight grey matter regions with between-scanner differences. In addition, noise and spatial inhomogeneities are variable across scanners: red arrows highlight such differences. B) Variability in T1-weighted images between (top) and within (bottom) scanners. There are few noticeable qualitative differences in the data between scanners. This is comparable to within-scanner data which is similarly consistent. Note: all data are shown in their respective native spaces.*

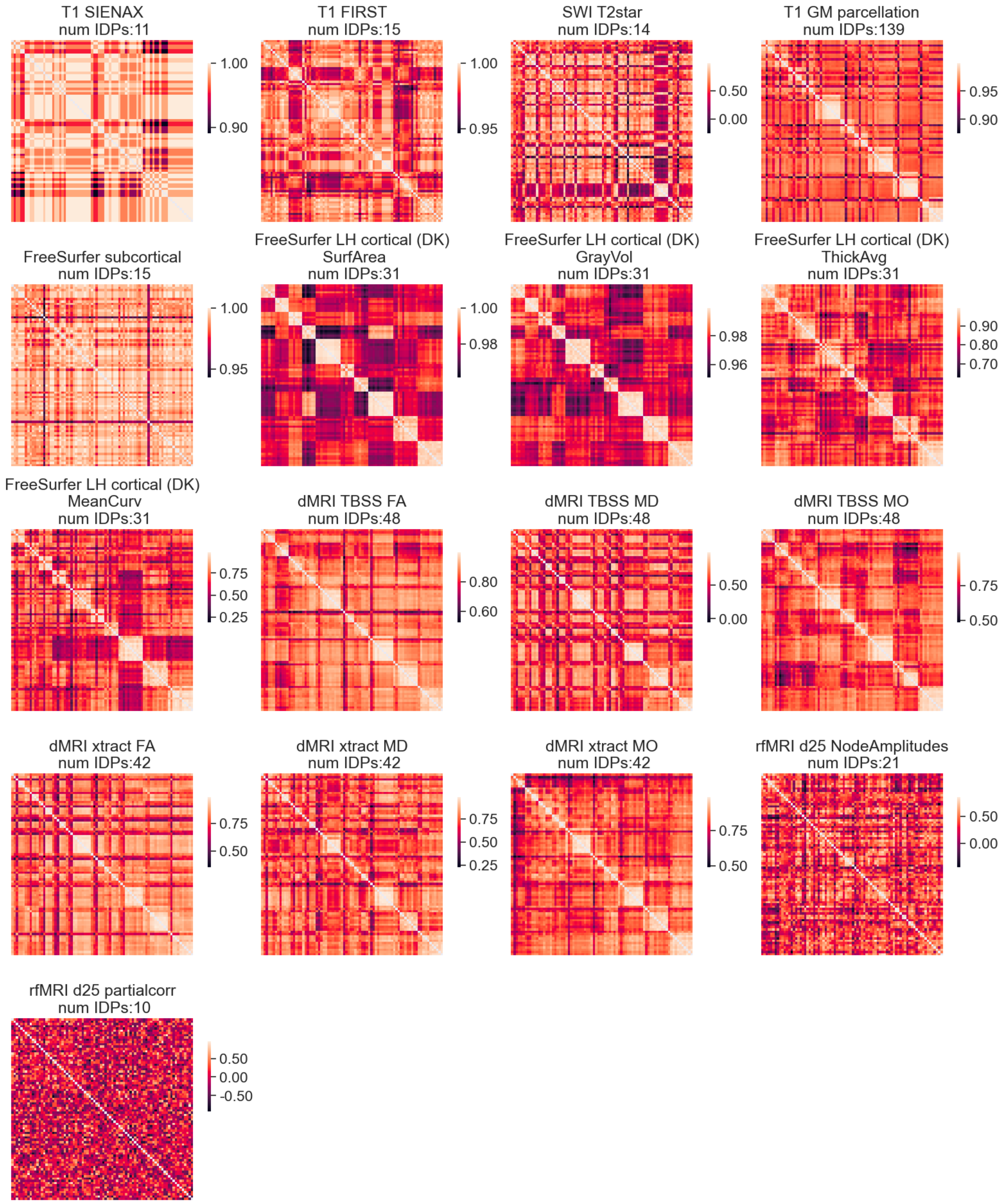

***Supplementary Figure 4*** *– Correlation (Spearman's rank) matrices $R_{ij}^{m}$ depicting the similarity of IDP categories ($M_{cat}$) between scanning sessions for all session pairs $(i, j)$. The number of IDPs in each IDP category is also reported. IDP categories include subcortical volumes, brain tissue volumes, subcortical T2*, cortical parcel volumes, dMRI regional and tract-wise microstructure (FA, MD, MO, L1, L2, L3), rfMRI functional connectivity node amplitude and edges.*

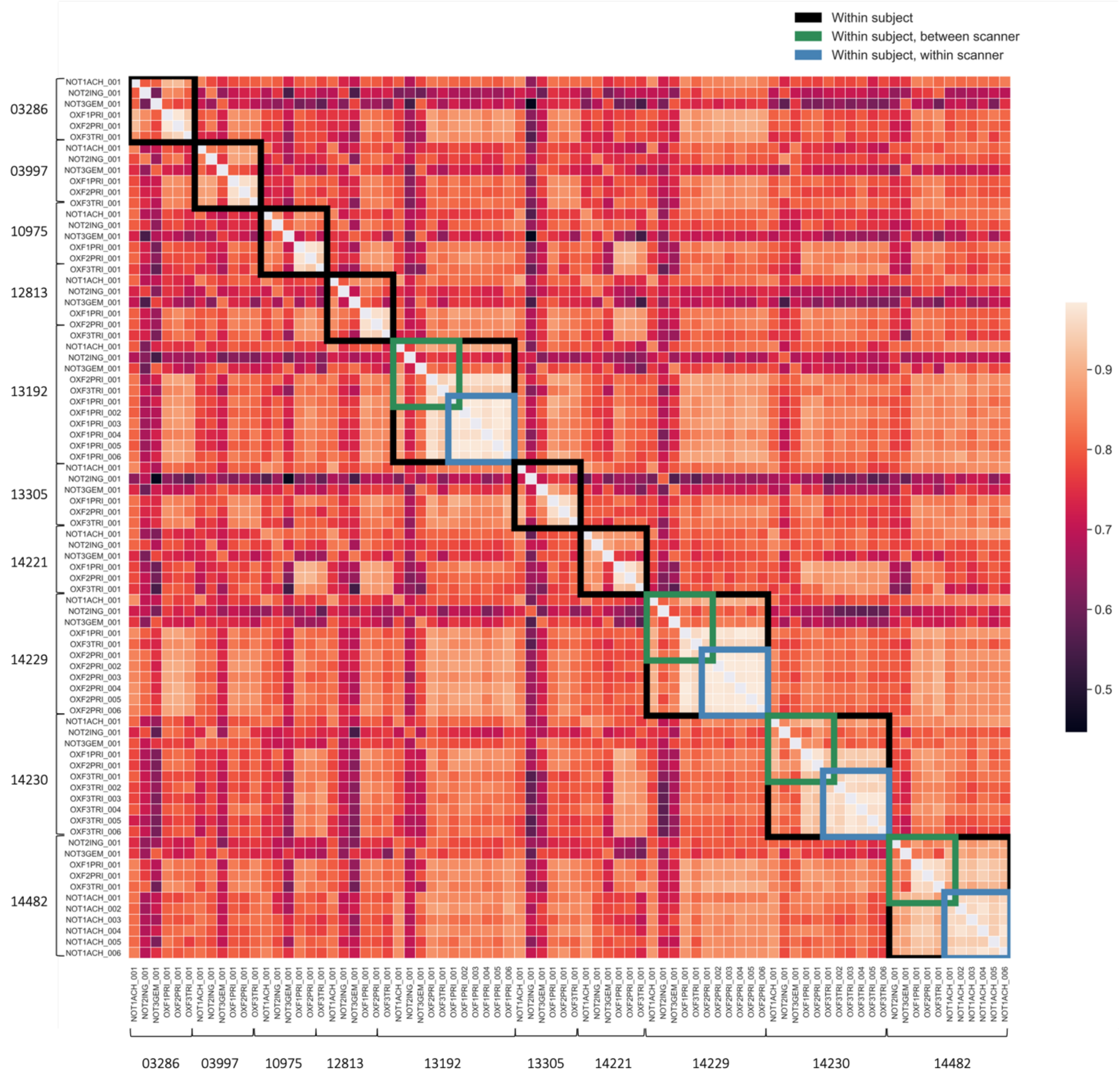

***Supplementary Figure 5** – The correlation (Spearman's rank) matrix $P_{ij}$ depicting the median similarity of IDPs between scanning sessions for all sessions $(i, j)$, calculated using Eq. 2.1 (Methods). Spearman's rank correlation is calculated for IDP categories ($M_{cat}$, presented in Supplementary Figure 4) and the median across categories plotted. Within-subject sessions are outlined in black. Within-scanner repeat sessions are outlined in blue. Between-scanner repeats are outlined in green.*

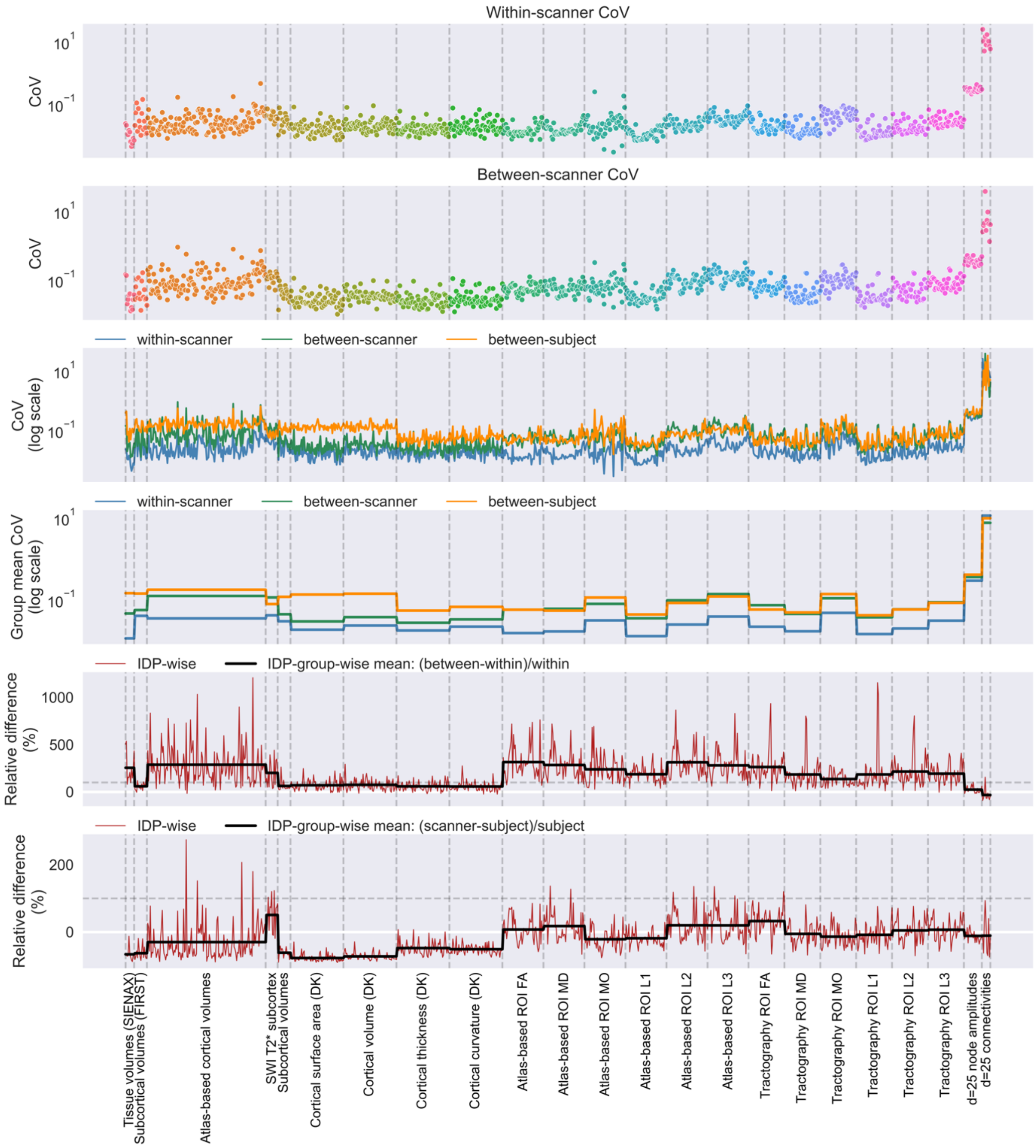

***Supplementary Figure 6*** *– As in Figure 6 but using only the four subjects with within-scanner repeats to calculate the within-scanner and between-scanner CoV. Top row: the IDP-wise CoVs across six within-scanner repeats, averaged across the four subjects with within-scanner repeats. Second row: the IDP-wise CoVs across six between-scanner repeats, averaged across the same four subjects. Third row: the within-scanner (blue), between-scanner (green) and between-subject-within-scanner (orange, reflecting biological variability) CoVs plotted on a log-scale. Fourth row: the IDP-group-wise mean of the CoVs plotted on a log scale for within-scanner (blue), between-scanner (green) and between-subject-within-scanner (orange) sessions. Fifth row: the IDP-wise (red) and IDP-group-wise (black) relative difference (between-within/within [scanner]) in CoVs. Bottom row: the IDP-group-wise relative difference between the scanner CoVs (within scanner, blue; between-scanner, green) and between-subject (biological) CoVs. The dashed horizontal line in rows five and six indicate relative difference of 100%.*

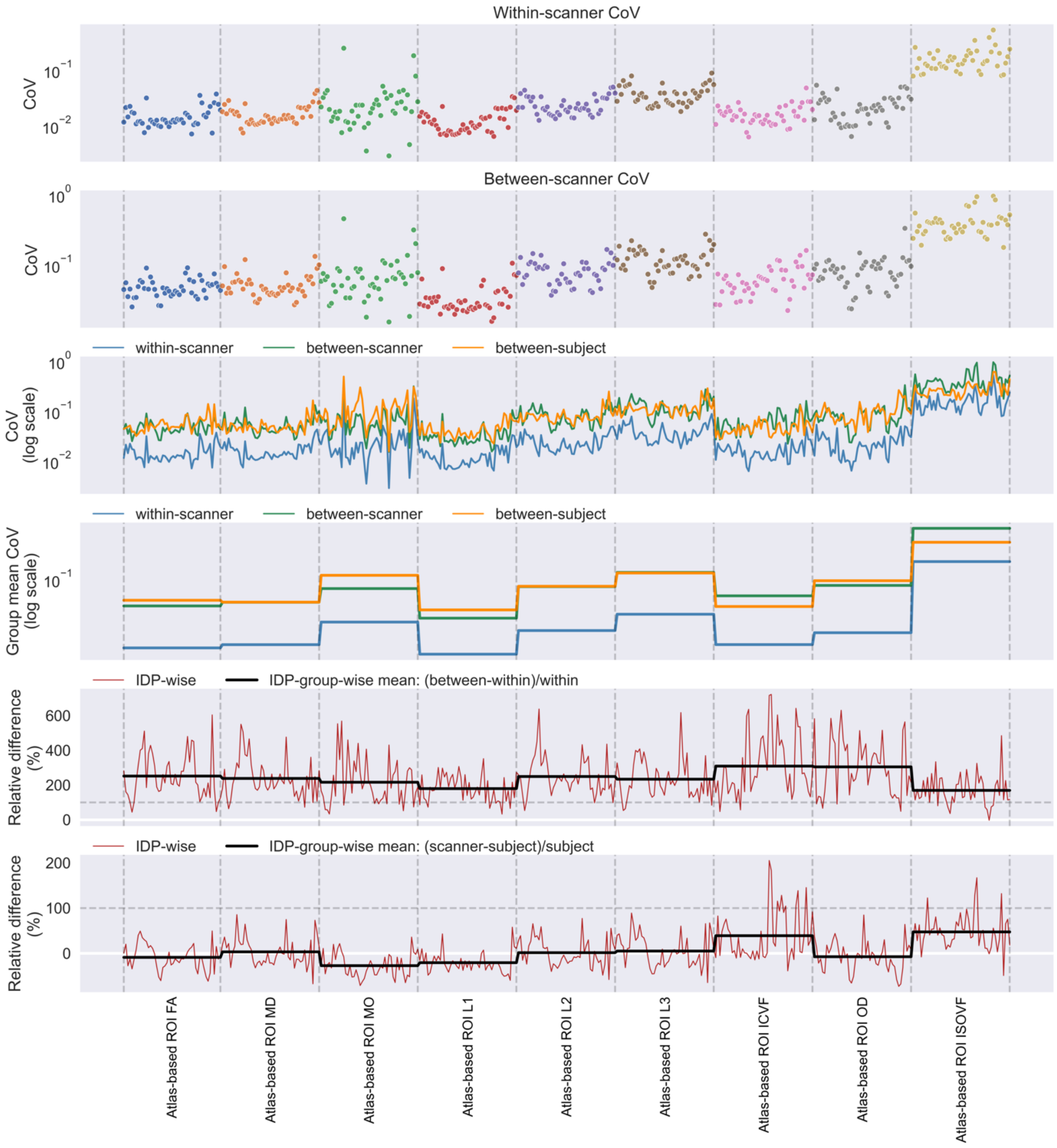

***Supplementary Figure 7** – The coefficient of variation of IDPs within/between scanner repeats for the atlas-based region wise diffusion metrics, including NODDI features. Top row: the IDP-wise CoVs across six within-scanner repeats, averaged across the four subjects with within-scanner repeats. Second row: the IDP-wise CoVs across six between-scanner repeats, averaged across all subjects. Third row: the within-scanner (blue), between-scanner (green) and between-subject-within-scanner (orange, reflecting biological variability) CoVs plotted on a log-scale. Fourth row: the IDP-group-wise mean of the CoVs (from the third row) plotted on a log scale for within-scanner (blue), between-scanner (green) and between-subject-within-scanner (orange) sessions. Fifth row: the IDP-wise (red) and IDP-group-wise (black) relative difference (between-within/within [scanner]) in CoVs. Bottom row: the IDP-group-wise relative difference in between-scanner CoVs (within scanner, blue; between-scanner, green) and between-subject (biological) CoVs. The dashed horizontal line in rows five and six indicate relative difference of 100%.*

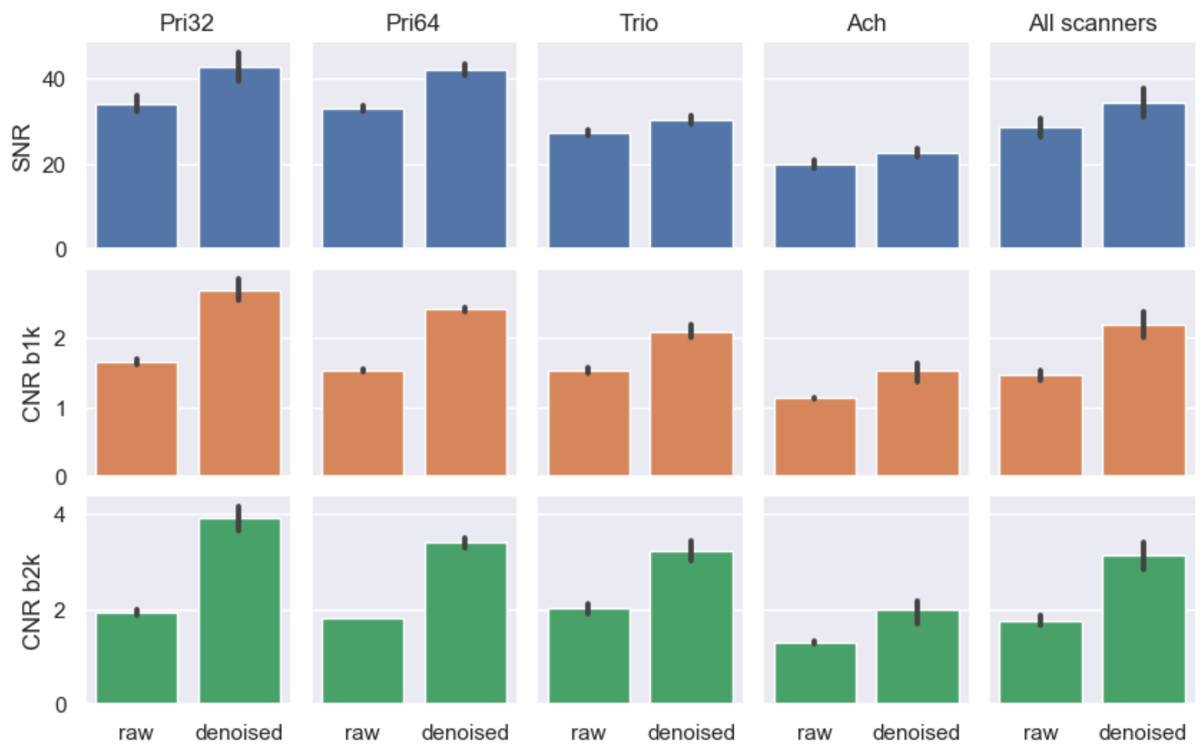

***Supplementary Figure 8** – Effect of diffusion MRI denoising on image quality metrics, including signal-to-noise ratio (SNR) and angular contrast-to-noise ratio (CNR) for different b values. Bars represent the mean (and standard deviation – error bars) of the reported IQM across subjects. Pri32: Siemens Prisma 32ch; Pri64: Siemens Prisma 64ch; Trio: Siemens Trio; Ach; Philips Achieva; All scanners: mean across scanners. SNR is reported for the b=0 s/mm$^2$ data; CNR b1k is the CNR for the b=1000 s/mm$^2$ data; CNR b2k is the CNR for the b=2000 s/mm$^2$ data.*
